# Fxr1 regulates sleep and synaptic homeostasis

**DOI:** 10.1101/709345

**Authors:** Jivan Khlghatyan, Alesya Evstratova, Lusine Bozoyan, Simon Chamberland, Aleksandra Marakhovskaia, Tiago Soares Silva, Katalin Toth, Valerie Mongrain, Jean-Martin Beaulieu

**Affiliations:** Department of Pharmacology & Toxicology, University of Toronto, Medical Sciences Building, Toronto, ON M5S 1A8, Canada; Department of Psychiatry and Neuroscience, Faculty of Medicine, Université Laval, Québec-City, QC G1J 2G3, Canada; Department of Neuroscience, Université de Montréal and Center for Advanced Research in Sleep Medicine, Hôpital du Sacré-Coeur de Montréal (CIUSSS-NIM), Montreal, QC H4J 1C5, Canada

**Author notes:** Now at: NYU Neuroscience Institute, New York University, Langone Medical Center, East River Science Park, New York, NY 10016, USA.

## Abstract

The fragile X autosomal homolog 1 (Fxr1) has been GWAS-associated to schizophrenia and insomnia but its contributions to brain functions are unclear. Homeostatic regulation of synaptic strength is essential for the maintenance of brain functions and engages both global and cell autonomous level processes. We used Crispr/Cas9-mediated somatic knockouts, overexpression, neuronal activity recordings and translatome sequencing, to examine the contribution of Fxr1 to cell-autonomous homeostatic synaptic scaling and global-level sleep homeostasis. Our findings indicate that Fxr1 is downregulated during scaling and sleep deprivation via a Gsk3β dependent mechanism. In both conditions, downregulation of Fxr1 is essential for the homeostatic modulation of synaptic strength. Furthermore, overexpression of Fxr1 during sleep deprivation results in altered EEG signatures and reverts changes of translatome profiles. These findings indicate that Fxr1 represents a shared signaling hub linking cell autonomous homeostatic plasticity and system level sleep homeostasis with potential implications for neuropsychiatric illnesses.

## Introduction

Regulation of synaptic strength is essential for the maintenance of proper brain functions and its disruption can contribute in circuit level imbalance of excitatory and inhibitory activity in neuropsychiatric illnesses such as Alzheimer disease, autism and schizophrenia ^1–3^. Homeostatic mechanisms, engaged in response to external conditions, are believed to contribute to this regulation both at a cell-autonomous and a global level ^4–9^. However, the relationship between these different spatial scales of regulation is unclear.

Synaptic scaling is a form of cell-autonomous homeostatic plasticity used by neurons to maintain net firing rates and is achieved by modulation of postsynaptic AMPA receptors ^10^. Prolonged inhibition of neuronal activity induces a multiplicative increase of miniature excitatory postsynaptic currents (mEPSCs) (upscaling) via increase of postsynaptic AMPA receptors. The opposite (downscaling) occurs as a result of prolonged neuronal activation ^11^.

At a global-level, homeostatic regulatory mechanisms are engaged during the sleep/wake cycle. Sleep pressure accumulates during wake and dissipates during sleep^12^. Studies suggest that neuronal activity correlates with sleep pressure by increasing during wake and decreasing during sleep ^6, 9, 13^. Moreover, both sleep pressure and neuronal activity can be further increased by prolonged wake and those changes can be reversed during a subsequent recovery period, indicating engagement of homeostatic mechanisms of regulation ^6, 9, 13^.

Some lines of evidence suggest that molecular and structural changes involved in downscaling also occur during sleep ^4, 14^. This raises the question of whether there are common homeostatic molecular regulators of neuronal activity involved in cell-autonomous synaptic scaling and global-level sleep homeostasis.

The fragile X mental retardation autosomal homolog 1 (Fxr1) might play a role both in sleep and homeostatic AMPA receptor regulation. Fxr1 is an RNA binding protein expressed in the brain and in neurons where it is localized in the cell bodies and dendrites in association with mRNAs and ribosomes ^15^. Variants of the *FXR1* locus are GWAS-identified risk factors for insomnia ^16^ and a schizophrenia-associated *FXR1* variant is linked to sleep duration ^17^. Interestingly, Fxr1 protein degradation is also regulated by Glycogen synthase kinase 3 beta (Gsk3β) ^18, 19^ and its brain expression is increased in response to lithium, a pharmacological agent inhibiting Gsk3 that is used for sleep regulation and the treatment of psychiatric disorders characterized by sleep disturbances, such as bipolar disorder ^20, 21^.

Along with Fxr2, Fxr1 is an autosomal paralogs of the fragile X mental retardation protein Fmrp, which is needed for the regulation of synaptic scaling by retinoic acid ^22, 23^. Fxr2, and Fmrp have been shown to regulate synaptic expression of the AMPA receptor GluA1 subunit in different brain regions ^24–26^. Fxr1 has been shown to regulate *de novo* synthesis of the GluA2 AMPA receptor subunit during long-lasting synaptic potentiation of hippocampal neurons ^27^. However, little else is known about the roles of Fxr1 in regulating brain functions.

The association of Fxr1 to insomnia led us to test whether it can be a regulator of synaptic and sleep homeostasis. Our results indicate that Fxr1 is engaged at a post-transcriptional level by sleep and synaptic homeostasis via a Gsk3β-dependent mechanism leading to its degradation. Furthermore, engagement of Fxr1 is essential and sufficient for the regulation of synaptic AMPA receptors in both systems, thus reveling a shared molecular underpinning between different scales of homeostatic regulation of synaptic strength.

## Results

### Fxr1 protein expression is reduced by homeostatic synaptic scaling

To evaluate the engagement of Fxr1 in homeostatic synaptic plasticity we induced homeostatic upscaling (TTX) or downscaling (BIC) in primary cortical cultures. Protein levels of all Fxr1 isoforms decreased during upscaling (TTX vs Veh) (Fig. 1a) with no changes during downscaling (BIC vs Veh) (Fig. 1b). Decreased protein levels during upscaling were not generalizable to other members of the fragile X family, with Fxr2 being marginally decreased (Fig. 1c) and Fmrp not being affected (Fig. 1d). Upscaling did not affect mRNAs levels of Fxr1 and Fxr2 (Fig. 1e,f) and induced an increase of Fmrp encoding mRNA (*Fmr1*) (Fig. 1g). This indicates that Fxr1 is regulated at a protein and not at mRNA level during upscaling. We have previously shown that Gsk3β can negatively regulate Fxr1 protein levels ^18^. Indeed, upscaling resulted in an increase of Gsk3α and β activity (decrease in inhibitory phosphorylation of Ser-21/9) and a decrease of Fxr1 protein (Fig. 1h-j). Conversely, inhibition of Gsk3α and β by lithium increased Fxr1 protein levels (Fig. 1h-j). Overall, this shows that Fxr1 protein is engaged specifically during upscaling and that this process involves Gsk3.

**Fig. 1.**
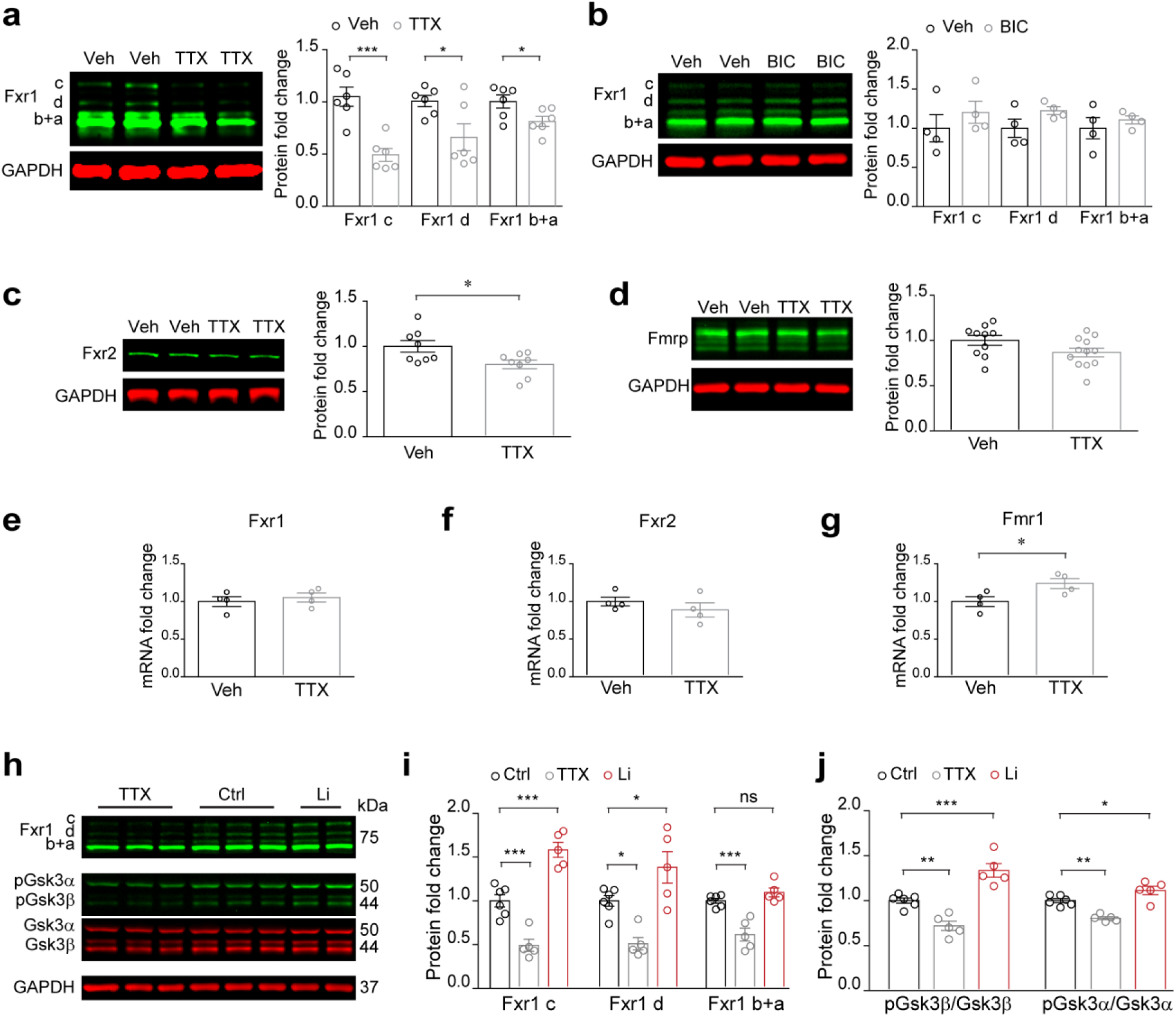
Fxr1 protein expression is decreased during homeostatic synaptic upscaling. Western blot analysis of Fxr1 during **a**, TTX induced upscaling (n=6 in each condition) and **b**, BIC induced downscaling (n=4 in each condition) of primary postnatal cortical cultures. Student’s T-test *p<0.05, ***p<0.001. Western blot analysis of **c**, Fxr2 (Veh n=8, TTX n=8) and **d**, Fmrp (Veh n=10, TTX n=12) during upscaling, Student’s T-test *p<0.05. RT-qPCR measurement of mRNA for **e**, Fxr1, **f**, Fxr2, **g**, Fmr1 during upscaling. n=4 in each condition, Student’s T-test *p<0.05. **h**, Western blot analysis for Fxr1, pGsk3α/β, Gsk3α/β and GAPDH in neuronal cultures treated with 1µM TTX (TTX) or 1mM LiCI (Li) or 1mM NaCI (Ctrl) for 48h. **i**, Expression of Fxr1 protein in TTX (n=5) and Li (n=5) conditions relative to Ctrl (n=6) condition. **j**, Expression of pGsk3β/Gsk3β or pGsk3α/Gsk3α in TTX (n=5) and Li (n=5) conditions relative to Ctrl (n=6) condition. One way ANOVA with Dunnett’s Multiple Comparison Test *p<0.05, **p<0.01, ***p<0.001. Error bars are mean ± SEM.

### Fxr1 suppresses the increase of surface GluA1 during synaptic upscaling

To investigate whether the decrease of Fxr1 protein is functionally contributing to upscaling, we verified whether an augmentation of Fxr1 could block upscaling. AMPA receptor expression is known to be modulated during homeostatic scaling ^28^. Fxr1 has been shown to negatively regulate translation of the GluA2 subunit of the AMPA receptor ^27^. Moreover, Fxr2 and Fmrp positively regulate synaptic GluA1 by stabilizing its mRNA or affecting membrane delivery respectively ^24^. Thus, we addressed whether Fxr1 can negatively regulate AMPA receptor expression during scaling.

Neuronal cultures were infected with high efficiency by AAV SYN GFP-Fxr1 (Fxr1 over) or AAV SYN GFP (Ctrl) (Fig. 2a, and Supplementary Fig. 1a,b) followed by measurement of GluA1 and GluA2 protein levels by Western blot during either up (TTX) or downscaling (BIC). Total GluA1 protein increased in Ctrl TTX condition as compared to Ctrl Veh (Fig. 2b). Surprisingly, this increase was exacerbated in the Fxr1 over condition following TTX treatment (Fxr1 over TTX vs Ctrl TTX) (Fig. 2b). No changes of total GluA1 protein was observed during downscaling (Fig. 2c). Total GluA2 protein did not change during upscaling (Supplementary Fig. 1d) or downscaling (Supplementary Fig. 1f). Consistently with results at the protein level, GluA1 mRNA increased in Ctrl TTX condition compared to Ctrl Veh and this increase was also further exacerbated in Fxr1 over TTX condition (Fxr1 over TTX vs Ctrl TTX) (Fig. 2d). This indicates that increase in GluA1 protein can be a result of a positive regulation by Fxr1 on an mRNA level.

**Fig. 2.**
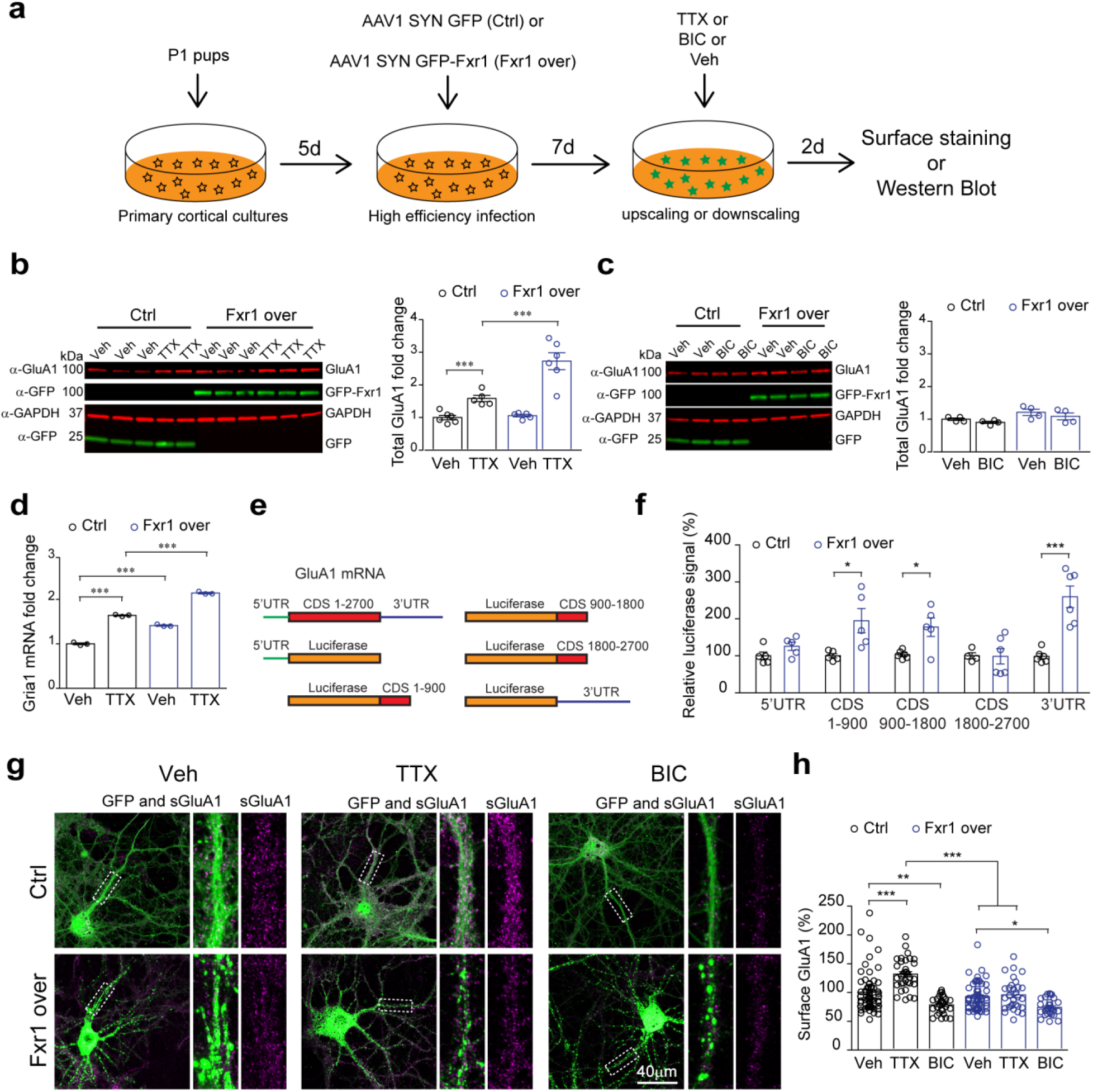
Fxr1 suppresses the increase of surface GluA1 during synaptic upscaling. **a**, Schematic of high-efficiency infection of neuronal cultures by AAV1 viruses followed by detection of AMPA receptor subunits. **b**, Western blot analysis of total GluA1 expression in Ctrl or Fxr1over condition during upscaling (Ctrl/Veh n=5, Ctrl/TTX n=5, Fxr1over/Veh n=6, Fxr1over/Veh n=6). One way Anova with Dunnett’s Multiple Comparison Test ***p<0.001. **c**, Western blot analysis of total GluA1 expression in Ctrl or Fxr1over condition during downscaling (Ctrl/Veh n=4, Ctrl/TTX n=4, Fxr1over/Veh n=4, Fxr1over/Veh n=4). **d**, RT-qPCR measurement of Gria1 mRNA in Ctrl or Fxr1over condition during upscaling (Ctrl/Veh n=3, Ctrl/TTX n=3, Fxr1over/Veh n=3, Fxr1over/Veh n=3). One way Anova with Bonferroni’s Multiple Comparison Test ***p<0.001. **e**, Schematic representation of tagging of Luciferase cDNA with 5’UTR, 1-900CDS, 900-1800CDS, 1800-2700CDS, and 3’UTR of GluA1 gene. **f**, Measurement of relative luciferase signal after co-transfection of different tagged luciferase constructs along with GFP (Ctrl) or GFP-Fxr1 (Fxr1 over) plasmids. n=5 in each condition, Student’s T-test *p<0.05, ***p<0.001. **g**, Immunostaining for surface GluA1 in GFP (Ctrl) or GFP-Fxr1 (Fxr1 over) infected cultures after treatment with Veh, TTX or BIC for 48h. **h**, Percentage of surface GluA1 relative to the mean of Ctrl/Veh condition (Ctrl condition: Veh n=63, TTX n=34, BIC n=29, Fxr1over condition: Veh n=56, TTX n=30, BIC n=29). One way Anova with Bonferroni’s Multiple Comparison Test *p<0.05, **p<0.01, ***p<0.001. Error bars are mean ± SEM.

To investigate whether Fxr1 can directly bind to GluA1 mRNA, we tagged a luciferase cDNA with 5’UTR, coding (CDS) or 3’UTR sequences from the GluA1 (*Gria1*) mRNA (Fig. 2e) and performed dual-luciferase assays. We detected an increase of luciferase signal from CDS1-900, CDS900-1800 and 3’UTR constructs in Fxr1 over condition compared to control (Fig. 2f) indicating a direct regulation to the potential binding sites. This shows that Fxr1 positively regulates levels of GluA1 during upscaling via direct binding to its mRNA.

Along with expression, surface levels of AMPA receptor are also changed during homeostatic scaling ^28^. Thus, we investigated the effect of Fxr1 on surface levels of AMPA receptors during up and downscaling. Surface expression of the GluA2 subunit did not change during upscaling or downscaling (Supplementary Fig. 1c,e). Surface expression of GluA1 increased during upscaling in Ctrl (Ctrl TTX vs Ctrl Veh) and this increase was abolished by Fxr1 overexpression (Fxr1 over TTX vs Fxr1 over Veh) (Fig. 2g,h). Decrease of surface GluA1 during downscaling was similar in both Ctrl and Fxr1 over conditions (Fig. 2g,h). No differences of surface GluA1 expression were observed between Ctrl Veh and Fxr1 over Veh conditions (Fig. 2g,h). This indicates that augmentation of Fxr1 expression can specifically block the increase of surface GluA1 during upscaling.

Overall, this shows that augmentation of Fxr1 blocks the increase of surface GluA1 during upscaling not via an inhibition of its expression but rather due to a negative regulation of GluA1 transport to the surface of the plasma membrane. Furthermore, this suggests that downregulation of Fxr1 can be required for the increase of surface GluA1 during upscaling.

### Decrease of Fxr1 is necessary and sufficient for the induction of multiplicative upscaling

Synaptic upscaling is a cell-autonomous process and it results in a multiplicative increase of miniature excitatory postsynaptic currents (mEPSCs) ^10, 11^. Fxr1 protein expression is engaged by upscaling via a mechanism that may involve Gsk3 and Fxr1 to regulate surface GluA1 during upscaling (Fig. 1 and 2). Thus, we aimed to understand whether the Gsk3β-Fxr1 signaling module can control synaptic currents in a cell-autonomous fashion during upscaling. To address this, we designed *Fxr1* gene targeting guide RNAs (gRNAs) and validated these using Neuro2A cells (Supplementary Fig. 2a-c). Then, the most efficient *Fxr1* targeting gRNA and a previously characterized *Gsk3b* targeting gRNA ^25^ were used to make Crispr/Cas9 constructs to target Fxr1 and Gsk3β in neuronal cultures. All control plasmids contained scrambled gRNAs. We generated Fxr1KO and Fxr1Ctrl (both tagged with mCherry), Gsk3KO and Gsk3Ctrl (both tagged with GFP) constructs and performed low-efficiency transfection of neuronal cultures (Supplementary Fig. 2d). Immunofluorescent staining revealed efficient single (Fxr1 or Gsk3β) or double (Gsk3β and Fxr1) knockouts of targeted genes (Supplementary Fig. 2e-h). An expression vector encoding GFP-Fxr1 was used for overexpression of Fxr1 (Fxr1over) and a vector expressing GFP alone was used as a control (Ctrl).

Whole-cell patch-clamp recordings showed a multiplicative increase in the mEPSC amplitude of control neurons following upscaling (Fig. 3a-d,o). Fxr1over or Gsk3KO prevented the increase of mEPSC amplitude induced by TTX (Fig. 3e,f,o). Fxr1KO resulted in an elevation of mEPSC amplitude in a multiplicative manner (Fig. 3g-j,o) to a level that was not further increased following activity blockade by TTX (Fig. 3g,o). Gsk3/Fxr1KO also induced multiplicative elevation of mEPSC amplitude (Fig. 3k-n,o) that was not further increased by TTX treatment similar to Fxr1KO (Fig. 3k,o). This shows that Fxr1 acts downstream of Gsk3β during upscaling. No changes in mEPSC frequency were observed in all the conditions (Fig. 3p).

**Fig. 3.**
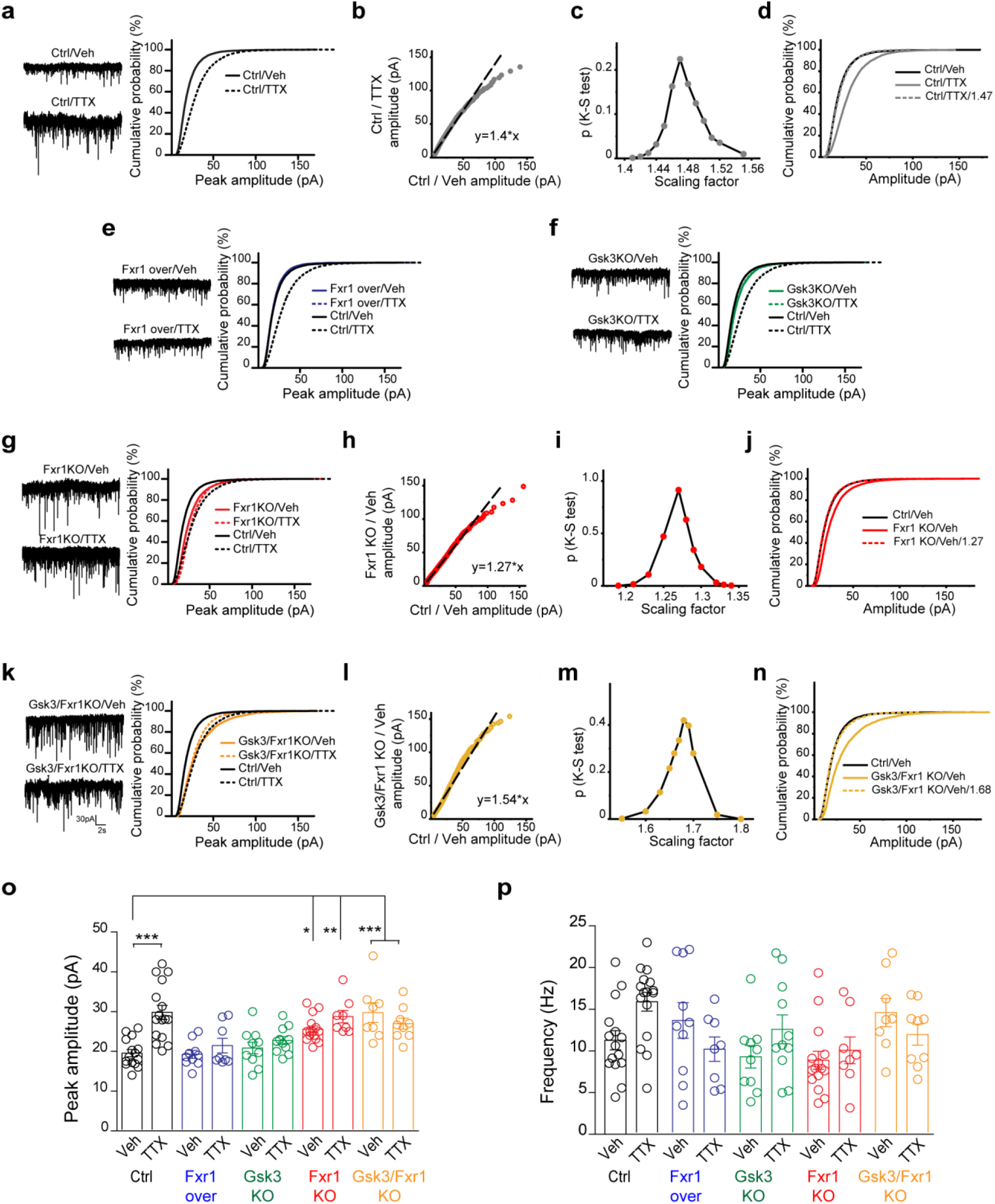
The decrease in Fxr1 expression is necessary and sufficient for induction of multiplicative upscaling. **a**, Cumulative probability plots of mEPSCs amplitude (500 events per cell) and representative examples of mEPSCs (left panel) recorded from cultured cortical control neurons after 48 hours of 1 µM TTX or Veh exposure (Ctrl/Veh n=16 and Ctrl/TTX n=17). **b**, A linear fit of Ctrl/TTX and Ctrl/Veh amplitudes. **c**, The degrees of overlap between Ctrl/TTX and Ctrl/Veh data were assessed using various scaling factors. The largest nonsignificant p-value was obtained with 1.47 scaling factor. **d**, Cumulative probability plots of the mEPSCs amplitude of Ctrl/Veh, Ctrl/TTX and Ctrl/TTX divided by scaling factor 1.47, which yielded the maximum overlap with Ctrl/Veh data. **e-g**, Cumulative probability plots of mEPSCs amplitude (500 events per cell) and representative examples of mEPSCs (left panel) recorded from cultured cortical neurons after 48 hours of 1 µM TTX or Veh exposure **e**, Fxr1P overexpressing neurons (Fxr1P over/Veh n=10 and Fxr1P over/TTX n=8), **f**, Gsk3 KO neurons (Gsk3KO/Veh n=10 and Gsk3KO/TTX n=11), **g**, Fxr1 KO neurons (Fxr1KO/Veh n=16 and Fxr1KO/TTX n=8). **h**, A linear fit of Fxr1 KO/Veh and Ctrl/Veh amplitudes. **i**, The degrees of overlap between Fxr1 KO/Veh and Ctrl/Veh data were assessed using various scaling factors. The largest non-significant p-value was obtained with 1.27 scaling factor. **j**, Cumulative probability plots of the mEPSCs amplitude of Ctrl/Veh, Fxr1 KO/Veh and Fxr1 KO/Veh divided by scaling factor 1.27, which yielded the maximum overlap with Ctrl/Veh data. **k**, Cumulative probability plots of mEPSCs amplitude (500 events per cell) and representative examples of mEPSCs (left panel) recorded from cultured cortical Gsk3 and Fxr1 KO neurons after 48 hours of 1 µM TTX or Veh exposure (Gsk3/Fxr1KO/Veh n=8 and Gsk3/Fxr1KO/TTX n=11). **l**, A linear fit of Gsk3/Fxr1 KO/Veh and Ctrl/Veh amplitudes. **m**, The degrees of overlap between Gsk3/Fxr1 KO/Veh and Ctrl/Veh data were assessed using various scaling factors. The largest non-significant p-value was obtained with 1.68 scaling factor. **n**, Cumulative probability plots of the mEPSCs amplitude of Ctrl/Veh, Gsk3/Fxr1 KO/Veh and Gsk3/Fxr1 KO/Veh divided by scaling factor 1.68, which yielded the maximum overlap with Ctrl/Veh data. **o**, mEPSC amplitude of cultured cortical neurons after 48 hours of 1 µM TTX or Veh exposure. One way Anova with Bonferroni’s Multiple Comparison Test *p<0.05, **p<0.01, ***p<0.001. **p**, mEPSC frequency of cultured cortical neurons after 48 hours of 1 µM TTX or Veh exposure. Error bars are mean ± SEM.

Overall, this indicates that during upscaling Gsk3β is activated and negatively regulates Fxr1 protein levels. In turn, the decrease in Fxr1 protein level is necessary and sufficient for the induction of homeostatic synaptic upscaling.

### Fxr1 modulates sleep duration and the response to sleep deprivation

To investigate the involvement of Fxr1 in the homeostatic regulation of neuronal activity on a global level, we examined whether Fxr1 is involved in sleep homeostasis. 48 hours of EEG and EMG recordings were performed in freely moving mice injected with AAV SYN GFP-Fxr1 (Fxr1over) or AAV SYN GFP (Ctrl) into the frontal cortex. Homeostatic sleep pressure accumulates during wake and is thus higher following enforced wakefulness or sleep deprivation (SD). We performed baseline (BL) recordings for 24h, followed by recordings during 6h of SD and 18h of recovery (REC). Measurement of the time spent in the wakefulness (WAKE), slow-wave sleep (SWS) and paradoxical sleep (PS) indicated that Fxr1over mice have a significantly different distribution of vigilant states over time in BL, an effect that became more pronounced during the REC period (Fig. 4a and Supplementary Fig. 3a). We computed the power spectra for WAKE, SWS, and PS during BL and REC (Fig. 4b). When normalized to the Ctrl, we noticed significant differences in the alpha frequency band (8 to 11.75 Hz) during wakefulness only in the REC period (Fig. 4c,d), which includes SD. Pronounced effects during REC can be indicative of involvement of Fxr1 in the response to elevated homeostatic sleep pressure.

**Fig. 4.**
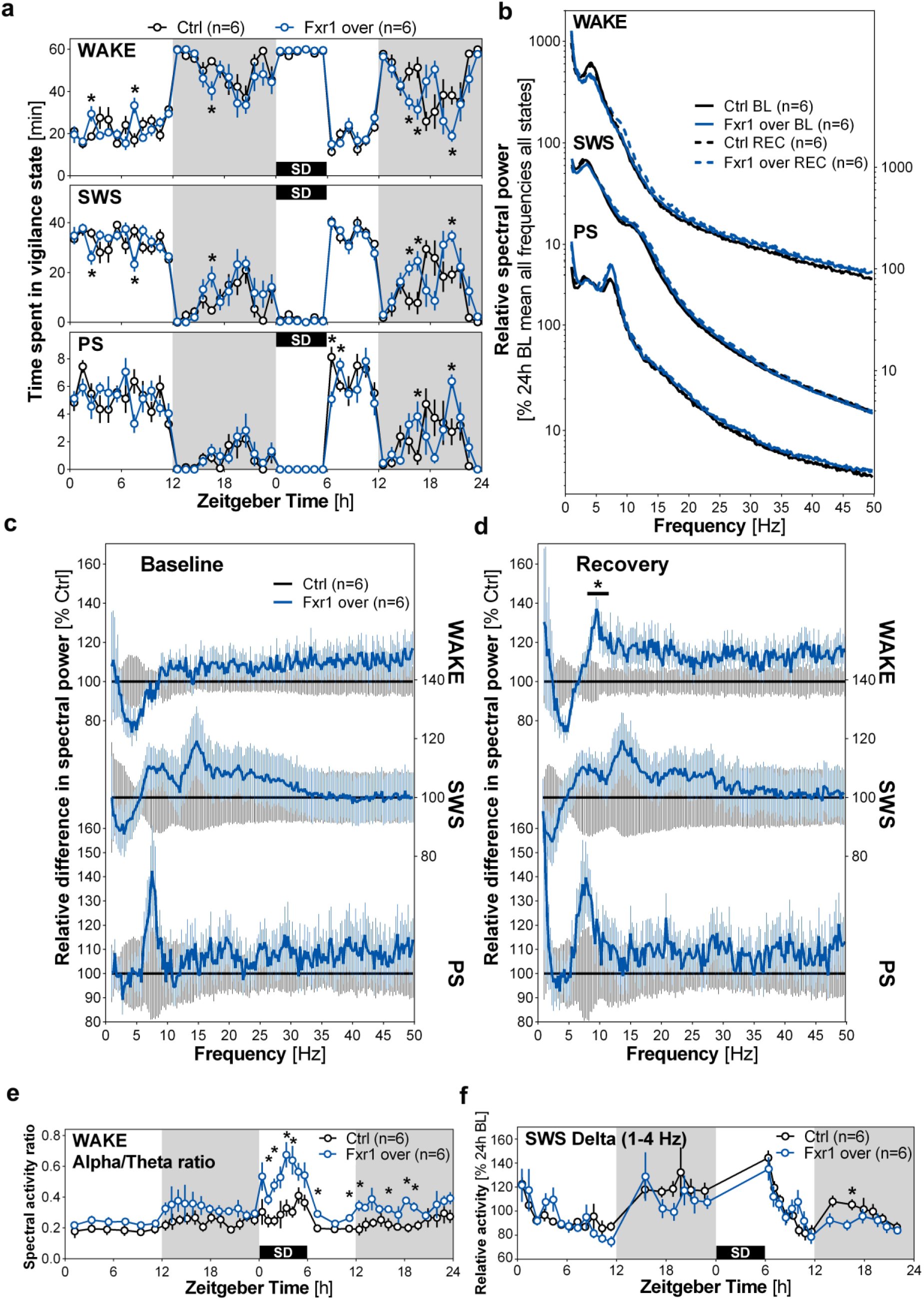
Fxr1 modulates sleep duration and the response to sleep deprivation. **a**, Hourly distribution of wakefulness (WAKE), slow-wave sleep (SWS) and paradoxical sleep (PS) during a 24-h baseline (BL) recording and a second 24-h starting with 6-h sleep deprivation (SD) (recovery: REC). Significant Group-by-Hour interactions were found for wakefulness during BL (F_23,230_ = 1.67) and REC (F_23,230_ = 2.83), for SWS during BL (F_23,230_ = 1.66) and REC (F_23,230_ = 2.89), and for PS during REC (F_16,160_ = 2.54). **b**, Power spectra for WAKE, SWS, and PS in Ctrl and Fxr1over mice computed between 0.75 and 50 Hz per 0.25-Hz for the full 24-h of BL and REC. **c** and **d**, The spectral activity of Fxr1over mice expressed relative to that of Ctrl mice for WAKE, SWS and PS during **c**, the 24-h BL and **d**, the 24-h REC. A significant difference between groups was found for the frequency band 8 to 11.75 Hz during wakefulness (t = 2.30). **e**, Time course of wakefulness spectral activity ratio between low alpha (8.5-10.5 Hz) and low theta (4-6 Hz) during BL and REC in Ctrl and Fxr1 over mice. A significant Group-by-Interval interaction was found for REC (F_22,220_ = 1.96). n=6 per group, Two-way ANOVA, Huynh-Feldt corrected, *p < 0.05. **f**, Time course of SWS delta activity during BL and REC in Ctrl and Fxr1over mice. A significant Group-by-Interval interaction was found during REC (F_13,130_ = 1.89). n=6 per group, Two way ANOVA, *p < 0.05.

SWS delta power increases with sleep pressure as found with SD, then it rapidly decreases during recovery sleep ^12^. Low alpha and high theta activity during WAKE were shown to importantly contribute to the homeostatic need for sleep ^29^. Moreover, decreased alpha/theta ratio was shown to associate with daytime sleepiness ^30^. First, we found an increase in alpha/theta ratio in Fxr1over mice compared to controls starting from the onset of SD (Fig. 4e and Supplementary Fig. 3b). Second, after SD, delta power in early dark phase was reduced in Fxr1over mice (Fig. 4f). These changes in SD effects in Fxr1over mice supports its contributions to sleep homeostasis.

### Fxr1 protein expression is engaged by sleep deprivation

Given that Fxr1 is involved in the regulation of EEG activity during SD, we thus investigated whether Fxr1 can be engaged by SD. Mice were divided into two groups, one group was sleep-deprived at the onset of the light phase (SD group), while the other was left undisturbed in the home cage (S group). Prefrontal cortex from both groups was dissected at the same time (Fig. 5a). SD induced an increase in synaptic p845 GluA1 (Fig. 5b) with no changes in GluA2 (Fig. 5c) as previously reported ^9^. Western blot analysis showed decreased levels of Fxr1 in the prefrontal cortex after SD (Fig. 5d). No changes of Fmrp protein were found during SD (Fig. 5e). SD induced a slight increase in mRNA of Fxr2 (Fig. 5g) with no changes of Fxr1 or Fmrp mRNAs (Fig. 5f,h). Overall, these results show that, in a manner similar to upscaling (Fig. 1), SD also results in a reduction of Fxr1 protein expression (Fig. 5).

**Fig. 5.**
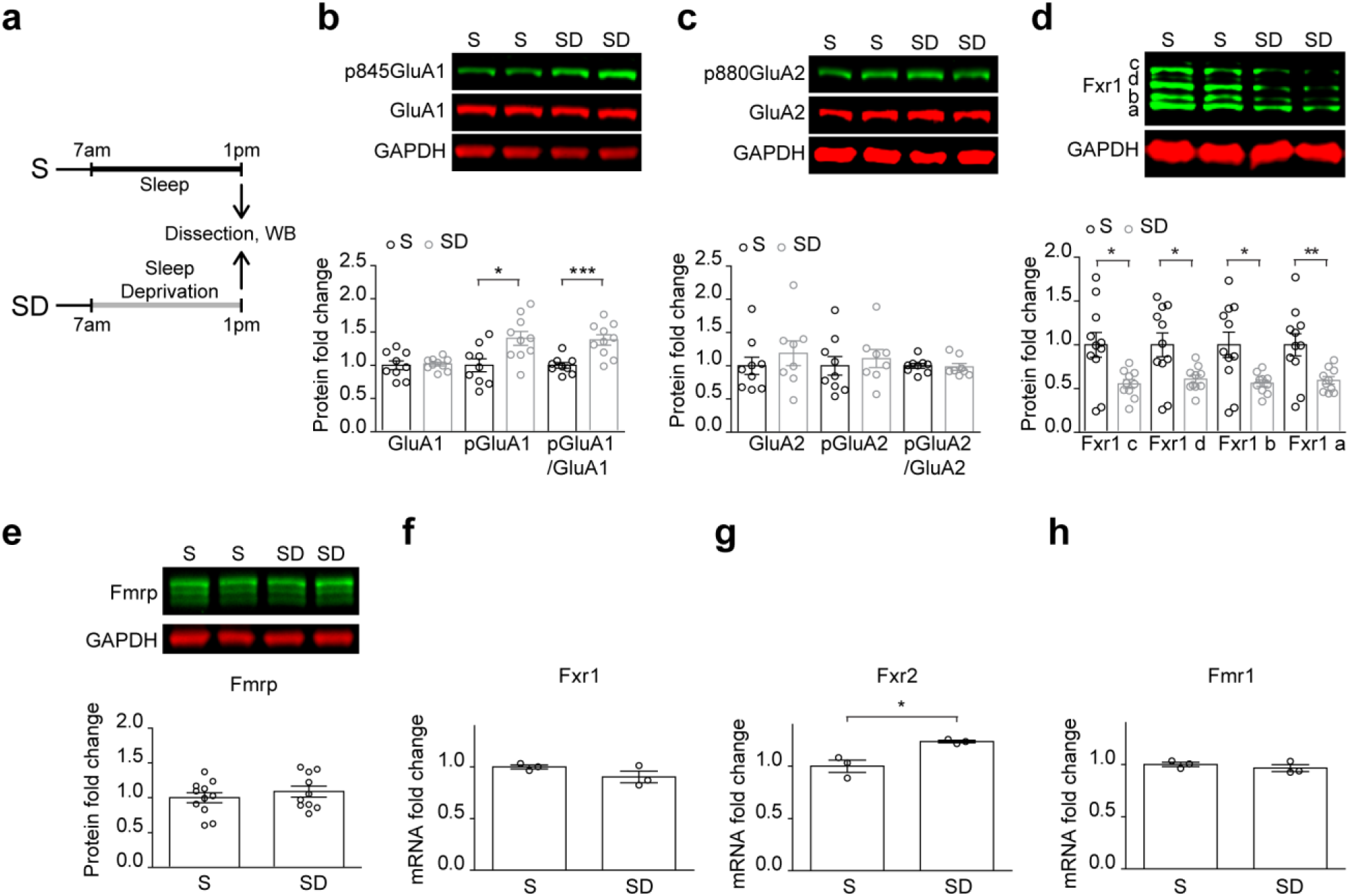
Fxr1 protein expression is decreased during sleep deprivation. **a**, Schematic representation of sleep deprivation experiments. **b-e**, Western blot analysis of **b**, p845GluA1 and GluA1 expression (S n=9, SD n=10), **c**, p880GluA2 and GluA2 expression (S n=9, SD n=8), **d**, Fxr1 expression (S n=11, SD n=10), and **e**, Fmrp expression (S n=11, SD n=10) in the prefrontal cortex of sleeping and sleep deprived mice. Student’s *t*-test p < 0.05, **p < 0.01, ***p < 0.001. **f-h**, RNAseq measurement of mRNA for **f**, Fxr1, **g**, Fxr2, **h**, Fmr1 during sleep deprivation. n=3 in each condition, Student’s T-test *p<0.05. Error bars are mean ± SEM.

### Fxr1 blocks increase of synaptic strength during sleep deprivation

It has been shown that SD results in an increase in mEPSCs ^13^. Fxr1 protein is engaged by synaptic upscaling (Fig. 1) and SD (Fig. 5) in a similar manner. Fxr1 negatively regulates synaptic GluA1 during upscaling (Fig. 2 and 3) and EEG signatures during SD (Fig. 4). Thus, we examined whether Fxr1 and its negative regulator Gsk3β can block increase of mEPSCs and synaptic GluA1 during SD. To address this question AAV SYN GFP-Fxr1 (Fxr1over), AAV Gsk3sgRNA/GFP + AAV SpCas9 (Gsk3sKO) ^25^, and AAV SYN GFP (Ctrl) were injected into the prefrontal cortex three weeks prior to SD (Fig. 6a). Whole-cell patch-clamp recordings on brain slices confirmed that SD increases mEPSC amplitude in the cortex of control mice (Fig. 6b,e), as previously shown ^13^. This effect was completely abolished in Fxr1over and Gsk3sKO conditions (Fig. 6c-e). No changes were observed in mEPSC frequency in all the conditions (Fig. 6f). Further characterization revealed a reduction of the rectification index indicating an increase of GluA1 containing calcium-permeable AMPA receptors in control mice (Fig. 6g,h). Conversely, this effect of SD was again abolished in Fxr1over or Gsk3sKO conditions (Fig. 6g,h). This indicates that augmentation of Fxr1 and inhibition of Gsk3β can prevent the increase of synaptic GluA1 during SD.

**Fig. 6.**
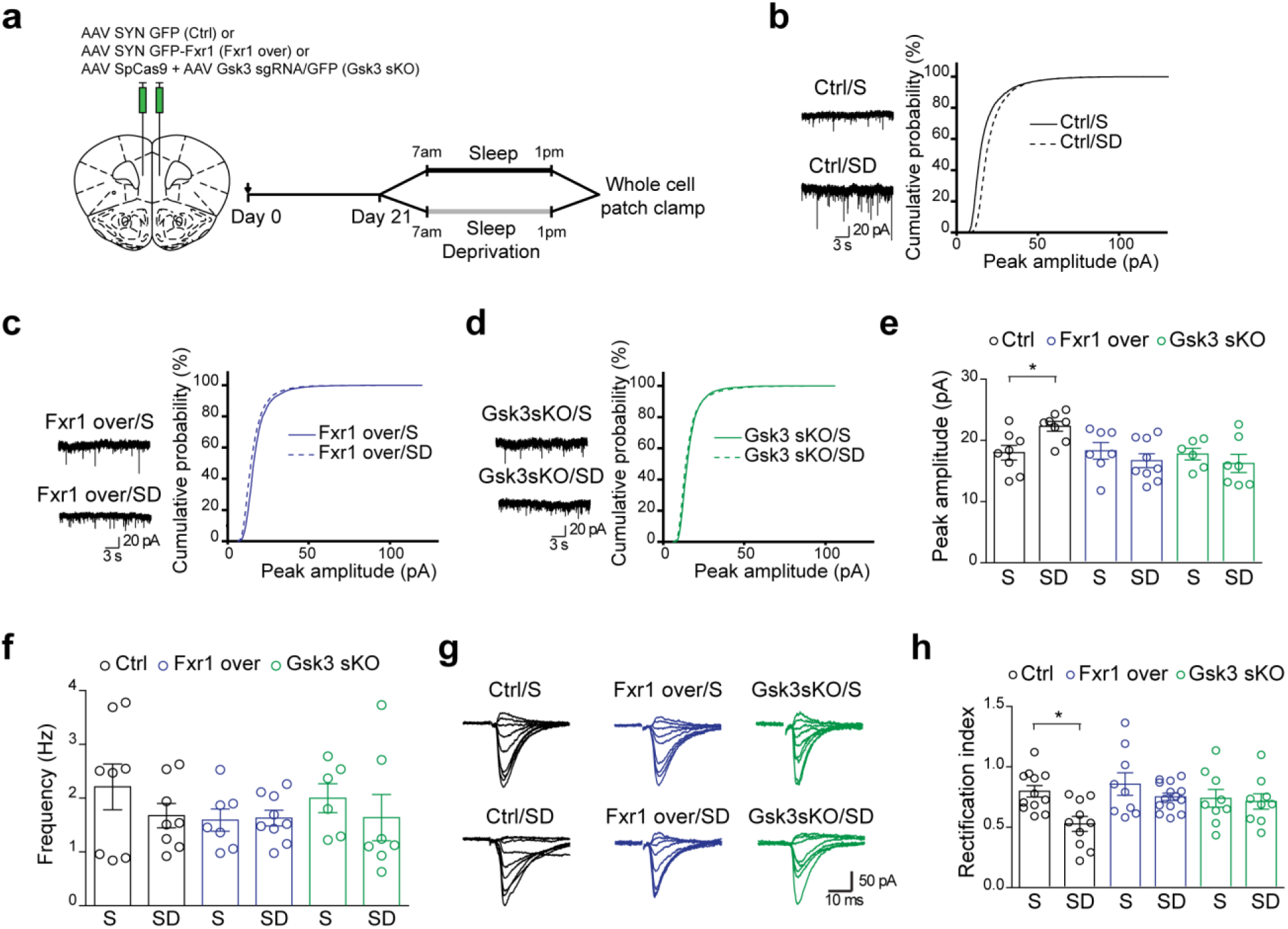
Fxr1 blocks increase of synaptic strength during sleep deprivation. **a**, Schematic representation of viral injection and sleep deprivation experiments. **b-d**, Cumulative probability plots of mEPSCs amplitude (500 events per cell) and representative examples of mEPSCs (left panel) recorded from brain slices of S and SD mice. **b**, Control neurons (Ctrl/S n=8, Ctrl/SD n=8), **c**, Fxr1 overexpressing (Fxr1 over/S n=7, Fxr1 over/SD n=9), and **d**, Gsk3sKO (Gsk3sKO/S n=6, Gsk3sKO/SD n=7). mEPSC **e**, amplitude and **f**, frequency of cortical neurons of S or SD mice. **g**, Representative examples of the current-voltage relationship of evoked EPSC amplitude recorded from S (top panel) and SD (bottom panel) mice. **h**, Summary bar graphs showing rectification index of control (S n=12, and SD n=10), Fxr1P overexpressing (S n=9, and SD n=14) and Gsk3 sKO (S n=9, and SD n=9) neurons. One way ANOVA with Bonferroni’s Multiple Comparison *p < 0.05. Error bars are mean ± SEM.

### Neuronal translatome regulation by Fxr1 during sleep deprivation

Fxr1 expression is engaged by SD (Fig. 5) and affects EEG signatures (Fig. 4) and synaptic activity (Fig. 6) during this process. We then aimed to investigate the impact of Fxr1 on SD on a molecular level. As an RNA binding protein, Fxr1 has a large number of targets ^31^, however, its neuronal targets have not been characterized. This makes it difficult to pinpoint direct molecular effectors of Fxr1 during SD. Thus, we used conditional RiboTag mediated RNA isolation ^32^ and translatome sequencing to find molecular signatures that are engaged by both sleep deprivation and Fxr1. This approach allows isolating ribosome-associated-RNAs only from cells that overexpress Fxr1. Moreover, unlike single cell or single nuclei techniques this approach provides a large amount of mRNA with a better signal/noise ratio ^33^ and preserves dendritic mRNA, which is important for local translation during plasticity.

RiboTag mice ^32^ were infected with AAV SYN GFP-Fxr1 + AAV SYN Cre (Fxr1over) or AAV SYN GFP + AAV SYN Cre (Ctr) viruses three weeks prior to sleep deprivation (Fig. 7a). This allowed activating RiboTag in the same neurons that overexpress Fxr1 (Fig. 7b,c). We found localization of GFP tagged Fxr1 along dendrites in close proximity with HA-tagged ribosomes (Fig. 7b) as previously reported for endogenous Fxr1 in cultured neurons ^15^.

**Fig. 7.**
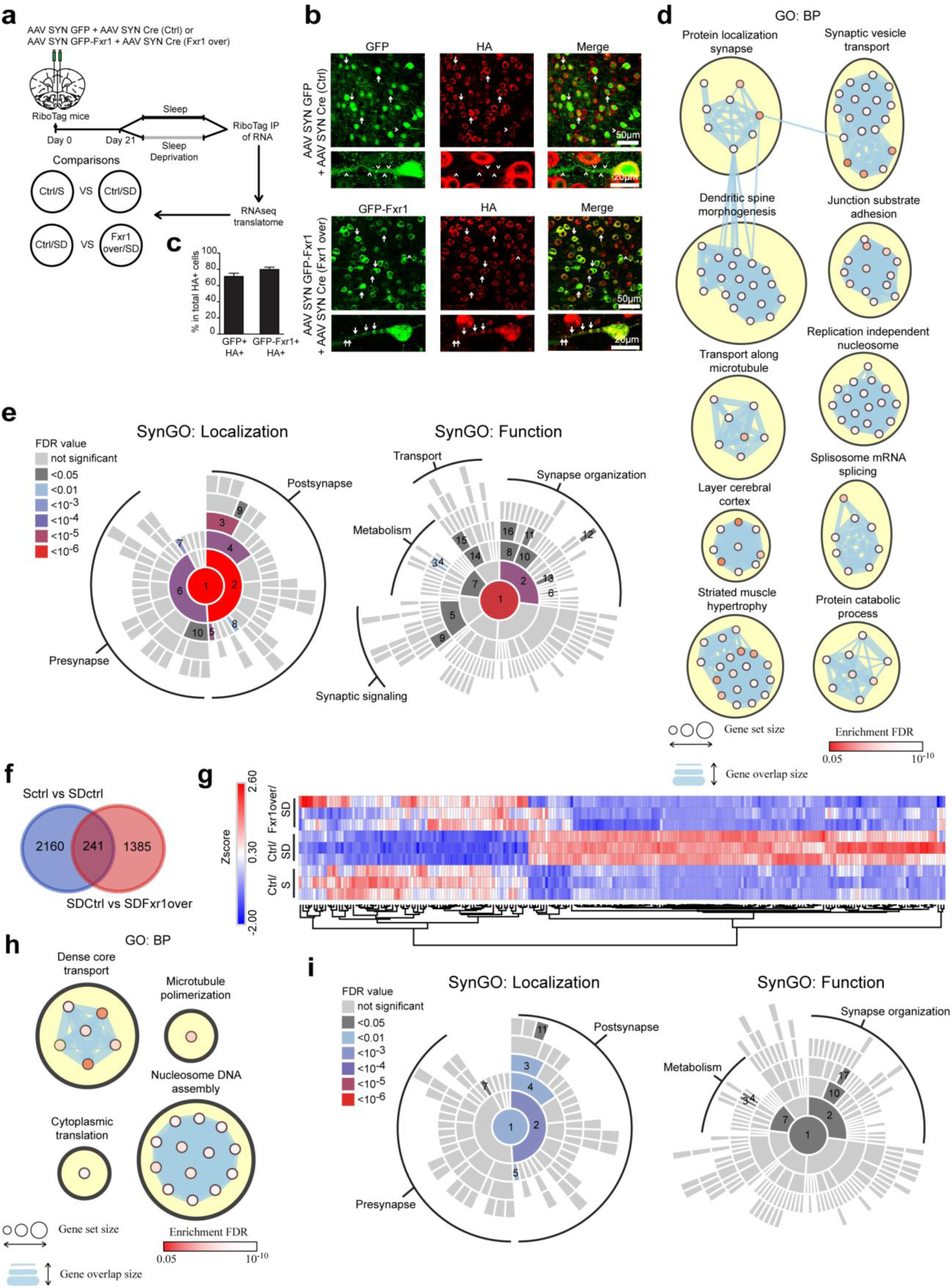
Neuronal translatome regulation by Fxr1 during sleep deprivation. **a**, Schematic representation of the experimental design. **b**, Immunostaining for GFP and HA in Ctrl and Fxr1over mouse brain slices. Arrows indicate presence and arrowheads indicate an absence of GFP tagged Fxr1 granules. **c**, Quantification of colocalization of GFP and HA labeled neurons. **d**, Enrichment of differentially expressed transcripts from Ctrl/S vs Ctrl/SD comparison in GO:BP. Top 10 clusters with the most number of enriched pathways (nodes) are shown. **e**, SynGO enrichment for synaptic localization and function of differentially expressed transcripts from Ctrl/S vs Ctrl/SD comparison. **f**, Venn diagram showing overlap of differentially expressed transcripts between Ctrl/S vs Ctrl/SD and Ctrl/SD vs Fxr1over/SD comparisons. **g**, Heat map showing transcripts that have bidirectional expression changes between Ctrl/S, Ctrl/SD, and Fxr1over/SD conditions. **h**, Enrichment of commonly affected transcripts between Ctrl/S vs Ctrl/SD and Ctrl/SD vs Fxr1over/SD comparisons in GO:BP. **i**, SynGO enrichment for synaptic localization and function of commonly affected transcripts between Ctrl/S vs Ctrl/SD and Ctrl/SD vs Fxr1over/SD comparisons. SynGO Localization: 1-Synapse, 2-Postsynapse, 3-Postsynaptic density, 4-Postsynaptic specialization, 5-Postsynaptic ribosome, 6-Presynapse, 7-Presynaptic ribosome, 8-Postsynaptic cytosol, 9-Postsynaptic density intracellular compartment, 10-Presynaptic active zone, 11-Integral component of postsynaptic density membrane. SynGO function: 1-Process in the synapse, 2-Synapse organization, 3-Protein translation at presynapse, 4-Protein translation at postsynapse, 5-Trans-synaptic signaling, 6-Synapse adhesion between pre- and post-synapse, 7-Metabolism, 8-Structural constituent of synapse, 9-Chemical synaptic transmission, 10-Synapse assembly, 11-Regulation of synapse assembly, 12-Regulation of modification of postsynaptic actin cytoskeleton, 13-Postsynaptic cytoskeleton organization, 14-Axo-dendritic transport, 15-Dendritic transport, 16-Structural constituent of postsynapse, 17-Postsynaptic specialization assembly.

After SD, RiboTag-associated-RNAs were isolated and subjected to sequencing followed by pairwise comparison (Fig. 7a). We identified 2401 unique differentially expressed transcripts (DETs) in Ctrl/S vs Ctrl/SD comparison (Supplementary Table 1). To identify biological dimensions based on DETs, we performed biological pathway enrichment (GO:BP) using gProfiler and clustering using EnrichmentMap and ClusterMaker in Cytoscape (Supplementary Table 4) ^34^. The top 10 largest clusters with the most significantly enriched pathways are shown in Fig. 7d. Five out of 10 clusters have relationship to the synapse (protein localization in synapse, synaptic vesicle transport, dendrite spine morphogenesis, transport along microtubule, junction substrate adhesion), thus we performed enrichment analysis using expert-curated and evidence-based synaptic gene ontology (SynGO) ^35^. This revealed that DETs engaged by SD (Ctrl/S vs Ctrl/SD) are highly enriched in synaptic localizations, particularly in post-synapse (Fig. 7e, Supplementary Table 6). Moreover, functional enrichment indicated involvement of those DETs in regulation of synapse organization, transport, metabolism and synaptic signaling (Fig. 7e, Supplementary Table 6).

Ctrl/SD vs Fxr1over/SD comparison identified 1626 unique DETs (Supplementary Table 2). We found 241 unique transcripts that are affected by both SD and Fxr1 during SD (Fig. 7f, Supplementary Table 3) (Ctrl/S vs Ctrl/SD overlap with Ctrl/SD vs Fxr1over/SD). For most transcripts (232), Fxr1 reversed changes induced by SD (e.g. upregulated by SD and downregulated by Fxr1 during SD) (Fig. 7g). These 232 transcripts were subjected to further analysis, GO:BP enrichment and clustering revealed 4 pathways (dense core vesicle transport, microtubule polymerization, cytoplasmic translation, nucleosome DNA assembly) (Fig. 7h, Supplementary Table 5). Localization analysis by SynGO identified enrichment in the postsynaptic compartments (Fig. 7i, Supplementary Table 7). SynGO functional analysis showed enrichment in synapse organization and metabolism (Fig. 7i, Supplementary Table 7). This shows that during SD, Fxr1 is involved in the regulation of multiple synaptic processes such as local translation and regulation of synaptic structure (Fig. 7e).

## Discussion

Homeostatic regulation of synaptic strength engages both cell autonomous and global levels mechanisms. Here we show that the insomnia GWAS-associated gene *Fxr1* encodes a common substrate involved in the regulation of synaptic strength during cell-autonomous synaptic scaling and global level sleep homeostasis. Fxr1 expression is engaged by SD and synaptic upscaling. Crispr/Cas9 and AAV mediated genetic manipulations of Fxr1 and its negative regulator Gsk3β controlled synaptic GluA1 and mEPSCs during upscaling and SD. Translatome sequencing indicated that Fxr1 is regulating local protein synthesis and synaptic structure related transcripts during SD. Based on our findings, we propose that during sleep deprivation and upscaling, Fxr1 protein level decreases due to its negative regulation by Gsk3β ^18, 19^. The decrease in Fxr1 protein induces broad alterations in synapse organization and metabolism, which subsequently result in an increase of synaptic GluA1 and postsynaptic excitatory activity.

We have shown that Fxr1 protein levels are only affected during upscaling and not downscaling (Fig. 1a,b). In line with this, augmentation of Fxr1 expression affected surface and total GluA1 expression only during upscaling (Fig. 2b-h) and not downscaling or basal conditions (Fig. 2b-h). Moreover, the increase of Fxr1 expression abolished upscaling (Fig. 3e,o) while decrease of Fxr1 was sufficient to induce upscaling (Fig. 3g-j,o). This shows that Fxr1 downregulation in response to Gsk3 is a necessary and sufficient mechanism inducing upscaling, thus underscoring the role of Fxr1 as a selective master regulator of this form of homeostatic plasticity.

Compared to other members of the fragile X family, Fxr1 expression is affected during upscaling (Fig. 1). Fmrp has also been shown to be involved in the homeostatic regulation of synaptic strength by retinoic acid ^23^. However, a lack of Fmrp expression abolishes upscaling in this system ^23^ and Fmrp protein levels are not affected by up scaling (Fig. 1). In contrast, Fxr1 is downregulated in response to TTX, Fxr1 KO is sufficient to induce upscaling (Fig. 3g-j,o), while Fxr1 over expression prevents this form of homeostatic plasticity. Differences in the contributions of Fxr1 and Fmrp can be partially explained by the mode of regulation of synaptic GluA1 subunit by these two proteins. Fmrp has been shown to facilitate GluA1 delivery to the membrane ^24^, while Fxr1 blocks this process during upscaling (Fig. 2). Similar to Fmrp, Fxr2 also positively regulates GluA1 albeit, via stabilization of its mRNA ^24^. Nevertheless, the role of Fxr2 in synaptic scaling has not been investigated. Overall, the engagement of Fxr1 by upscaling and its role in regulating synaptic GluA1 functionally dissociates Fxr1 from the other two members of the fragile X family.

Furthermore, the regulation of synaptic GluA1 appears to be specifically engaged in response to external conditions, thus supporting its contribution to homeostatic responses. Augmentation of Fxr1 expression had no effect on mEPSCs *in vitro* (between Fxr1over/Veh and Ctrl/Veh conditions) (Fig. 3e,o,p) and *in vivo* (between Fxr1over/S and Ctrl/S conditions) (Fig. 6e,f,h). Augmentation of Fxr1 affected mEPSCs only during upscaling (Fxr1over/TTX vs Ctrl/TTX) (Fig. 3e,o) or sleep deprivation (Fxr1over/SD vs Ctrl/SD) (Fig. 6e,h). This indicates activity-dependent engagement of Fxr1 in regulation of GluA1 both during upscaling and sleep deprivation.

Fxr1 expression was not affected by downscaling (Fig. 1b) and augmentation of Fxr1 expression did not affect downscaling (Fig. 2b,c,e) or EEG activity during sleep (Fig. 4). Fxr1 affected EEG activity during WAKE in the course of SD and after (Fig. 4d,e), during SWS in the recovery dark phase (Fig. 4f) and upscaling (Fig. 2 and 3). This indicates the possible engagement of distinct molecular mechanisms of scaling based on sleep/wake state. Indeed, downscaling mechanisms appear to be engaged during sleep ^4^. Moreover, upscaling mechanisms occurring as a result of visual deprivation are only engaged in wake and are suppressed by sleep ^5^. Our study thus indicates that some molecular regulators of homeostatic upscaling, such as Fxr1, may also contribute to the dynamics of sleep homeostasis.

The mechanisms by which Fxr1 is engaged both by SD and upscaling necessitate Gsk3β activity and a downregulation of Fxr1protein levels. Fxr1 has been shown to be phosphorylated by Gsk3β and targeted for degradation following it ubiquitination ^18, 19^. However, this regulation involves the priming of Fxr1 by other kinases and may thus integrate information from several upstream signaling pathways^18^. Similarly, Fxr1 also regulates the translation, trafficking and stability of several mRNA^31, 36, 37^. In the context of SD in prefrontal cortex neurons, our results indicate that the translation of multiple transcripts involved in neuronal processes including synapse organization and metabolism are regulated by Fxr1. Taken together, these observations suggest that the Gsk3β-Fxr1 module may constitute a shared signaling hub involved in the homeostatic regulation of synaptic strength both at the cell autonomous and global levels in response to environmental-allostatic loads such as SD ^38, 39^.

The contribution of Fxr1 to the homeostatic regulation of synaptic strength can be important for several human disorders. Indeed, variants in the Fxr1 locus have been GWAS-associated to insomnia ^16^ and mental illness including schizophrenia and bipolar disorder ^40, 41^. Fxr1 is also differentially expressed in the brain of people with schizophrenia ^42^, while a schizophrenia-associated SNP in *Fxr1* has been linked to self-reported sleep duration ^17^.

The involvement of Fxr1 in sleep homeostasis is supported by its engagement by SD and the involvement of Fxr1 in the regulation of neuronal activity at a cellular (Fig. 5 and 6) and EEG levels during SD, as well as the regulation of duration of vigilant states (Fig. 4). The interaction between *Fxr1* and *Gsk3b* have also been reported to affects anxiety-related behaviors in mice ^25^ and self-reported emotional stability in humans ^18^. Furthermore, expression of Fxr1 can be regulated by lithium and other mood stabilizer drugs as a consequence of Gsk3 inhibition ^18^. Neuropsychiatric disorders and lithium have been associated with regulation/dysregulation of sleep ^20, 43^ and homeostatic plasticity ^44, 45^. Further investigations of the Gsk3β-Fxr1 signaling module should clarify if it also constitutes a molecular link between mental illnesses, sleep homeostasis and cell-autonomous homeostatic plasticity.

## Acknowledgments

Authors acknowledge Chloé Provost and Julien Dufort-Gervais for technical help with AAV injection/EEG implantation surgery. JMB is Canada Research Chair in Molecular Psychiatry. VM is Canada Research Chair in Sleep Molecular Physiology. This work was supported by grants from Canada Institutes of Health Research (CIHR, PJT-148568) to JMB, and salary awards from CIHR and Fonds de recherche du Québec - Santé (FRQS) to VM. JMB is NARSAD independent investigator and One-Mind Rising Star awardee.

## Author contributions

JMB conceived the study. JMB and JK designed the experiments. JMB, JK, AE, and VM wrote the manuscript. JK performed design and testing of CRISPR/Cas9 *in vitro* and *in vivo*, stereotaxic injections, protein expression analysis *in vitro* and *in vivo*, primary neuronal culture preparation, drug treatment, and receptor surface staining and quantification, sleep deprivation, Ribo-Tag IP and RNA extraction and RNAseq data analysis. AE and SC performed whole-cell patch-clamp recordings and data analysis. AM performed CRISPR/Cas9 KO experiments with puromycin selection followed by detection of Fxr1 expression. LB performed cloning and luciferase assay. VM performed EEG recordings and analyses. VM, AM, and TSS participated in mouse sleep deprivation experiments. VM and KT provided technical, financial and intellectual support.

## Data and materials availability

RNAseq data and analysis will be deposited to GEO. For additional materials please contact the lead author.

## Online Methods

### Experimental model and subject details

All experiments conducted in this study are approved by the Université Laval, the CIUSSS-NIM and the University of Toronto Institutional Animal Care Committees in line with guidelines from the Canadian Council on Animal Care.

### Experimental animals

For primary cultures, postnatal day 0-1 (P0-P1) pups of C57BL/6J mice were used. For stereotaxic injection followed by ribosome immunoprecipitation (RiboTag-IP) 3-4 months old RiboTag mice ^32^ were used. For all other experiments C57BL/6J male (Jackson Laboratory, Bar Harbor ME) mice were used. Littermates were housed 3-4 per cage in a humidity-controlled room at 23°C on a 12 h light-dark cycle with ad libitum access to food and water. At the time of experiment mice were 3-4 months old and weighed approximately 25-30g. All animals were used in scientific experiments for the first time. This includes no previous exposures to pharmacological substances or altered diets.

### Primary cultures

Primary cortical cultures were prepared from P0-P1 C57BL/6J mouse pups as described ^46^. Briefly, neurons were dissociated from cortices of P0-1 pups and seeded on poly-D-lysine coated coverslips (neuVitro, Vancouver, WA) in 24 well plates. For western blot analysis and electrophysiology, neurons were seeded at the density of 5×10^5^ cells per well. For surface expression analysis neurons were seeded at the density of 1.5×10^5^ cells per well. Cultures were grown at 37°C and 5% CO2 in Neurobasal plus medium, supplemented with B27 plus, Glutamax (Thermo Fisher Scientific, Waltham, MA) and penicillin/streptomycin mix for a total of 13-14 days.

### DNA constructs

To knockout (KO) *Fxr1* gene 20-nt target sequences in exons of this gene were selected using online CRISPR design tool (http://crispr.mit.edu/) to minimize off-target activity. For in vitro testing, guide oligonucleotides (targeting *Fxr1*) were cloned into pX330 (pX330-U6-Chimeric_BB-CBh-hSpCas9 was a gift from Feng Zhang (Addgene # 42230)) ^47^ all in one vector by single-step cloning using BbsI restriction sites ^48^. For primary neuronal culture transfection previously characterized Gsk3b gRNA oligonucleotide ^25^ was cloned into pX458 (pSpCas9(BB)-2A-GFP (PX458) was a gift from Feng Zhang (Addgene plasmid # 48138)) ^48^ vector by single-step cloning using BbsI restriction sites to generate Gsk3 KO construct. pX458 vector was used as a control (Gsk3 Ctrl construct). To generate Fxr1 Ctrl construct first GFP from pX458 vector was changed to mCherry. mCherry was PCR amplified by Forward (mCherryF) 5’-AATAATGAATTCGGCAGTGGAGAGGGCAGAGGAAGTCTGCTAACATGCGGTGACGTCGAGGAGAATCCTGGCCCAGTGAGCAAGGGCGAGGAGGATAACA-3’ and Reverse (mCherryR) 5’-AATAATGAATTCTTACTTGTACAGCTCGTCCATGC-3’ primers and inserted into the pX458 vector using EcoRI restriction sites. To generate the Fxr1 KO construct the most active *Fxr1* targeting guide (Fxr1 gRNA2) oligonucleotide was cloned into Fxr1 Ctrl vector by single-step cloning using BbsI restriction sites. For in vitro KO of *Fxr1* in Fig. S3F guide was cloned into pX459 vector (pSpCas9(BB)-2A-Puro (PX459) V2.0 was a gift from Feng Zhang (Addgene plasmid # 62988)) ^48^. Sequences of all constructs were verified.

### AAV viral particle preparation

AAV serotype 1 SYN GFP-Fxr1 viral particles were produced by the University of North Carolina (UNC) Vector core facility. AAV serotype 5 SpCas9, Gsk3sgRNA/GFP, SYN GFP-Fxr1 are previously characterized ^25^ and are also produced by the University of North Carolina (UNC) Vector core facility. AAV serotype 1 GFP (AAV1 SYN GFP), AAV serotype 5 GFP (AAV5 SYN GFP) and AAV serotype 5 Cre (AAV5 SYN Cre) were purchased from Addgene.

### Cell line culture and transfection

Neuro-2A (N2A) cells were grown in high glucose DMEM containing 10% FBS, penicillin/streptomycin and L-glutamine (HyClone-GE Healthcare, Logan, UT). Cells were maintained at 37°C in 5% CO2 atmosphere and transfected using Lipofectamine 2000 (Thermo Fisher Scientific, Waltham, MA) according to the manufacturer’s protocols.

To test the activity of Fxr1 sgRNA by western blot (Fig. S3F), 50–70% confluent N2A cells were transfected with all in one px459 based constructs (pX459 vectors with guide targeting *Fxr1*). To select only transfected cells, 48 hours after transfection cells were incubated with 3µM puromycin for 72 hours followed by 48 hours incubation without puromycin. Cells were washed and lysed on day 7 after transfection.

### Genomic DNA extraction and SURVEYOR assay

For functional testing of sgRNAs, 50–70% confluent N2A cells were transfected with all in one pX330 based constructs (pX330 vectors with guides targeting *Fxr1*). Cells transfected with pX330 only served as a negative control. Cells were lysed 48 h after transfection by tail buffer (Tris pH=8.0 0.1M, NaCl 0.2M, EDTA 5mM, SDS 0.4% and proteinase K 0.2mg/ml), and DNA was precipitated using isopropanol followed by centrifugation (13000g 15min). DNA was resuspended in TE Buffer (10 mM Tris pH 8.0, 0.1 mM EDTA) and used for downstream analysis. Functional testing of individual sgRNAs was performed by SURVEYOR nuclease assay (Transgenomics, Omaha, NE) using PCR primers listed below. Band intensity quantification was performed as described ^48^.

*PCR primers used in the SURVEYOR assay:*

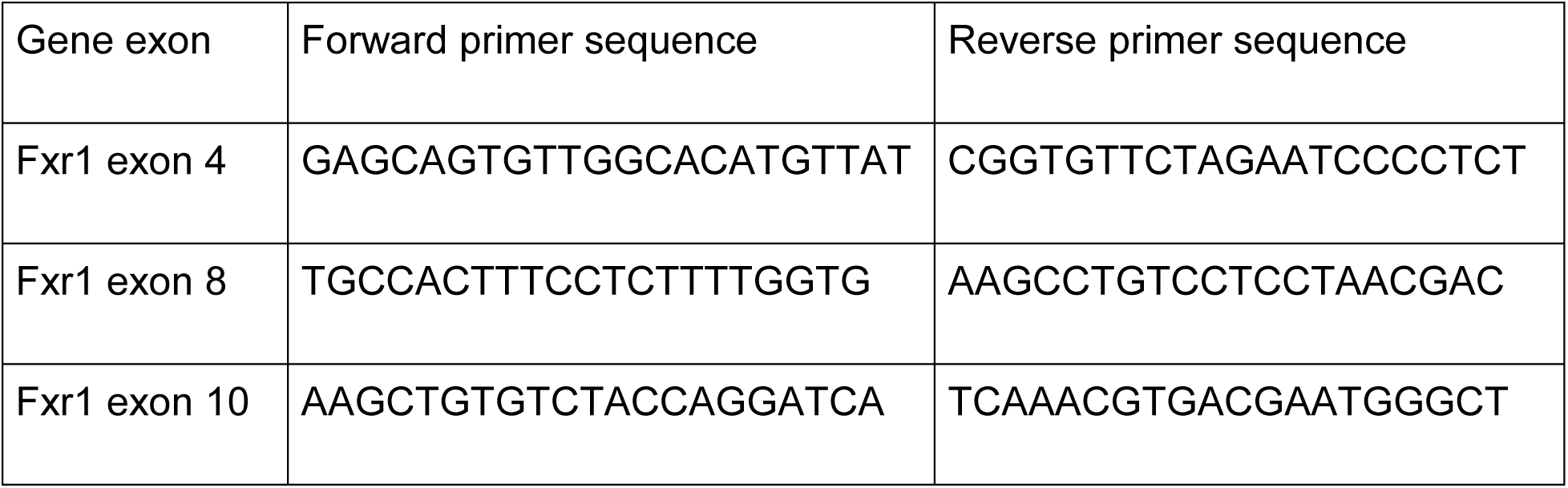

### Primary cortical culture transfection and infection

Cultures were transfected at DIV7 (with Gsk3 Ctrl, Gsk3 KO, Fxr1 Ctrl, Fxr1 KO constructs) and at DIV10 (with Fxr1 over construct) using DNA In-Neuro transfection (MTI GlobalStem, Gaithersburg, MA) reagent according to manufacturer’s protocols. Cultures were infected at DIV5 with AAV1 SYN GFP or AAV1 SYN GFP-Fxr1. Viruses were added to the cultures at the titer of 2×10^4^ viral genomes (Vg)/per neuron. To induce upscaling or downscaling cell were incubated in the presence of 1µM TTX or 50µM BIC (Alomone Labs, Jerusalem, Israel) or vehicle respectively from DIV12 to DIV14. All experiments were performed at DIV14.

### Stereotaxic injections

3 weeks before the sleep deprivation and electrophysiology recordings, bilateral injection of the virus was made in the prefrontal cortex. Mice were anesthetized with a preparation of ketamine 10mg/ml and xylazine 1mg/ml (0.1ml/10 g, i.p.). The animal was placed in a stereotaxic frame, and the skull surface was exposed. Two holes were drilled at injection sites and 1ul of virus (AAV GFP-Fxr1 4.4×10^12^vg/ml or AAV GFP 4.5×10^12^vg/ml or 1:1 AAV mixture: AAV SpCas9 2.6×10^12^vg/ml and AAV Gsk3sgRNA/GFP 5.4×10^12^vg/ml or AAV SpCas9 and AAV GFP 4.5×10^12^vg/ml, AAV SYN GFP-Fxr1 4.4×10^12^vg/ml and AAV SYN Cre 5.5×10^12^vg/ml or AAV SYN GFP 4.5×10^12^vg/ml and AAV SYN Cre 5.5×10^12^vg/ml) was injected using nanoliter injector with microsyringe pump controller (WPI) at the speed of 4nl per second. Following coordinates were used: anterior-posterior (AP), +2.4 mm anterior to bregma; mediolateral (ML), ±0.5 mm; dorsoventral (DV), 1.7 mm below the surface of the brain. All measures were taken before, during, and after surgery to minimize animal pain and discomfort.

### Surgeries for EEG recordings

The surgery combined viral delivery to the frontal cortex with electrode implantation for EEG/EMG recording that has been performed as described previously ^49–51^. Mice were anesthetized (Ketamine/Xylazine 120/10 mg/kg, intraperitoneal injection) and placed in a stereotaxic frame. A 28 gauge cannula was positioned for injection to the right frontal cortex (1.5 mm anterior to bregma, 1.5 mm lateral to midline and 1.5 mm below the skull surface) and delivered 1 μl of AAV-GFP (4.7×1012 vg/ml) or of AAV-Fxr1-GFP (4.4×1012 vg/ml) at an injection speed of 0.025 μl/min. The cannula was left in place for 5 min after the injection and then slowly removed. Two gold-plated screws (EEG electrodes; diameter 1.1 mm) were screwed through the skull over the right cerebral hemisphere: the first above the AAV injection site and the second above the posterior cortex (1 mm anterior to lambda, 1.5 mm lateral to the midline). An additional screw serving as a reference electrode was implanted on the right hemisphere (0.7 mm posterior to bregma, 2.6 mm lateral to the midline) together with three anchor screws implanted on the left hemisphere. Two gold wires were implanted between neck muscles and served as EMG electrodes. Electrodes were soldered to a connector and fixed to the skull with dental cement. Four days after surgery, mice were connected to a swivel relay and habituated to the cabling condition until recording.

### EEG recording and analyses

EEG/EMG recording and analyses were performed similarly to previously described ^49–51^. Briefly, signals were amplified with Lamont amplifiers, and sampled at 256 Hz and filtered using the software Stellate Harmonie (Natus, San Carlos, CA). A bipolar montage was used for the visual identification of vigilance states (wakefulness, slow-wave sleep [SWS], paradoxical sleep [PS]) on 4-sec epochs and the time spent in each state was computed per hour as well as per 12-h light and dark periods. Spectral analysis was computed using fast Fourier transform on artifact-free epochs of the anterior/frontal EEG signal (referenced to the reference electrode) to calculate spectral power per 0.25-Hz bins from 0.75 Hz to 50 Hz and per frequency bands (i.e., SWS delta [1-4 Hz], wakefulness low theta [4-6 Hz] and low alpha [8.5-10.5 Hz]). Power spectra were normalized relative to the total power of all states and also as a function of spectral activity in the control group. The spectral activity of frequency bands was averaged for intervals containing an equal number of SWS or wakefulness epochs and expressed relative to 24-h baseline mean as described before ^49, 51, 52^. More precisely, to take into account the distribution of wakefulness and SWS, SWS delta activity was average for 12 equal intervals during the baseline light period, 6 equal intervals during the dark periods and 8 equal intervals during the 6-h light period following SD; whereas wakefulness low theta and low alpha activity were averaged for 6 equal intervals during the baseline light period, 12 equal intervals during dark periods, 8 equal intervals during SD and 3 equal intervals during the 6-h light period following SD. A frequency band activity ratio was computed by dividing low alpha by low theta absolute activity separately for each interval. The distribution of vigilance states and the time course of activity in frequency bands have been plotted according to Zeitgeber time (Zeitgeber time 0 = Lights ON; Zeitgeber time 12 = Lights OFF).

### Statistical analyses for EEG recordings

Hourly distribution of vigilance state duration and time course of spectral activity in frequency bands have been compared between groups (Ctrl vs. Fxr1 over) using two-way repeated-measure analyses of variances (ANOVAs). Significance levels were adjusted for repeated measures using the Huynh-Feldt correction and significant interactions were decomposed using planned comparisons. Time spent in vigilance states computed per 12-h periods and spectral activity in the alpha band during wakefulness was compared between groups using t-tests. The threshold for statistical significance was set to 0.05 and data are presented as the mean and standard error of the mean.

### Ex-vivo electrophysiological recordings and drugs

Mice were taken for experiments 6 hours after the onset of the light phase with or without sleep deprivation.

### Acute slice preparation

Mice were killed by rapid cervical dislocation. Cortical slices (300 μm) were prepared from mice (3 weeks after injection of viruses) using a vibrating-blade microtome (Leica Biosystem, Wetzlar, Germany). Slices were prepared using ice-cold artificial cerebrospinal fluid (ACSF) containing: NaCl 87mM, NaHCO3 25mM, KCl 2.5mM, NaH2PO4 1.25mM, MgCl2 7mM, CaCl2 0.5mM, glucose 25mM and sucrose 75mM. Right after sectioning, slices were placed in oxygenated ACSF at 32°C for 30 min, transferred to extracellular ACSF and maintained at room temperature prior to experiments. All recordings were performed with extracellular ACSF containing: NaCl 124mM, NaHCO3 25mM, KCl 2.5mM, MgCl2 1.5mM, CaCl2 2.5mM and glucose 10mM, equilibrated with 95% O2/5% CO2, pH7.4, maintained at 31–33°C and perfused at a rate of 2-3 mL/min.

### Electrophysiology

Whole-cell current-clamp and voltage-clamp recordings were made with glass electrodes (4–6.5 MΩ) filled with a solution containing: K-gluconate 120mM, KCl 20mM, MgCl2 2mM, EGTA 0.6mM, MgATP 2mM, NaGTP 0.3mM, Hepes 10mM, phosphocreatine 7mM or Cs-gluconate 100mM, NaCl 8mM, MgCl2 5mM, EGTA 0.6mM, MgATP 2mM, NaGTP 0.3mM, Hepes 10mM, phosphocreatine 7mM, QX-314 1, spermine 0.1mM (Cs-gluconate-based solution was used to investigate I-V relationships of evoked EPSCs,). Pyramidal neurons expressing either GFP (green) or mCherry (red) were visually identified in acute slices (mPFC layer III-V) and cortical cultures using a fluorescence microscope. Electrophysiological recordings were made using a Multi Clamp 700A amplifier (Axon Instruments, Union City, CA), operating under current-clamp and voltage-clamp mode. Data were filtered at 4 kHz, and acquired using pClamp 10 software (Molecular Devices, Sunnyvale, CA). All recordings were done at a holding potential −70 mV, except sleep deprivation experiments, where holding potential was −60 mV. For the I-V curve experiments holding potential was varied from 100 mV to 60 mV. The uncompensated series resistance was monitored by the delivery of −10 mV steps throughout the experiment, only recordings with less than 15% change were analyzed.

### Drugs

10 μM CNQX, 50 μM AP5 and 10 μM bicuculline methiodide (Sigma-Aldrich, Oakville, Canada) were dissolved in extracellular ACSF and applied through the perfusion system (at least five minutes before recordings). 0.5 µM TTX were always present in the ACSF if recordings were done in cultures.

### Analysis of electrophysiological recordings

Synaptic events were analyzed using pClamp 10 software within at least 3 minutes of recordings, individual events were detected using an automatic template search. Templates were created using the average of at least 10 events aligned by the rising of their slopes. The peak amplitude of evoked EPSCs (eEPSCs) was measured for an averaged response (5 trials). Paired-pulse ratio was calculated as average for 15-20 trials. Rectification index (RI) was calculated, as a ratio of *I*–*V* slopes, RI= *s*2/*s*1 ^53, 54^. First, we calculated slope 1 (s1) using linear regression to *AMPA currents* recorded at holding potential ≤ 0 mV, as well as an AMPAR reversal potential, *E* _rev_. Next, we estimated slope 2 (s2) using a linear fit of *I*–*V* data recorded at positive holding potentials and constrained to intersect the x-axis at *E*_rev_. This method allows taking into account variations of AMPA reversal potential between recordings.

### Immunofluorescent staining

Mice were euthanized 3 weeks after viral delivery by a lethal dose of ketamine/xylazine and perfused with phosphate buffer saline (PBS) followed by 4% paraformaldehyde (PFA). Brains were incubated in 4% PFA 24h at 4°C. Fixed tissue was sectioned using vibratome (Leica, VT1000S). Next, 40 μm sections were blocked and permeabilized with a permeabilization solution containing 10% normal goat serum (NGS) and 0.5% Triton X-100 (Sigma) in PBS for 2 h. Sections were incubated with primary antibodies diluted in permeabilization solution overnight at 4 °C. After three washes in PBS, samples were incubated with secondary antibodies for 2h at room temperature. After washing with PBS three times, sections were mounted using DAKO mounting medium (DAKO, Mississauga, Canada) and visualized with a confocal microscope (Zeiss LSM 700, Zen 2011 Software, Oberkochen, Germany).

For immunofluorescent staining of primary neurons, cells were fixed at DIV14 with 4% paraformaldehyde (PFA) 4% sucrose mixture for 7 min at room temperature (RT). After washing three times with PBS, cells were permeabilized with 0.1% Triton X-100 in PBS for 10 min at RT. In the case of surface staining for GluA1 and GluA2, no permeabilization was performed. Cells were washed once with PBS and blocked by 10% serum in PBS for 1 hour. Cells were incubated with primary antibodies in 1% serum in PBS overnight at 4 °C. After washing three times with PBS, cells were incubated with secondary antibodies for 2 h at RT. Finally, coverslips were mounted using DAKO mounting medium and imaged using Zeiss LSM 880. Images were processed using the Zen 2011 (Zeiss, Oberkochen, Germany). Quantifications of colocalization were performed manually using ImageJ (National Institute of Health (NIH), Bethesda, MD). For quantification of surface GluA1 and GluA2, 2-3 dendrites (ranging from 50µm to 150µm) per cell were delineated and mean signal intensity was measured using ImageJ (National Institute of Health (NIH), Bethesda, MD). Following primary antibodies were used: Mouse anti-GluA1 (1:1000, Millipore MAB2263), mouse anti-GluA2 (1:1000, Millipore MAB397), rabbit anti-Fxr1 (1:1000, Abcam 129089), mouse anti-Gsk3β (1:500, Abcam 93926).

Secondary antibodies: Alexa Fluor 405, 568 or 647 (Life Technologies/ Thermo Fisher Scientific, Waltham, MA, 1:1000).

### Western blot

Mice were killed by cervical dislocation, after which the heads of animals were immediately cooled by immersion in liquid nitrogen for 6 s. The medial part of the prefrontal cortex was rapidly dissected out (within 30 s) on an ice-cold surface and frozen in liquid nitrogen before protein extraction. Tissue samples were homogenized in boiling 1% SDS solution and boiled for 5 min before measurement of protein concentration. Neuro2A cells and primary cortical cultures were lysed in lysis buffer containing: 50mM Tris-HCl, 150mM NaCl, 5mM EDTA, Protease inhibitor cocktail, 1% SDS, 0.5% Na-deoxycholate, 1% NP-40, 10mM NaFluoride, 25mM βglycerophosphate, 10mM Na Orthovanadate (Sigma-Aldrich, Oakville, Canada). Lysates were centrifuged 10000g for 30 min and supernatants were collected. Protein concentration was measured by using a DC-protein assay (Bio-Rad, Hercules, CA). Protein extracts were separated on precast 10% SDS/PAGE Tris-glycine gels (Thermo Fisher Scientific, Waltham, MA) and transferred to nitrocellulose membranes. Blots were immunostained overnight at 4 °C with primary antibodies. Immune complexes were revealed using appropriate IR dye-labeled secondary antibodies from Li-Cor Biotechnology (Lincoln, NE). Quantitative analyses of fluorescent IR dye signal were carried out using an Odyssey Imager and software (Licor Biotechnology, Lincoln, NE). For quantification, GAPDH (Actin in case of Neuro2A cells) was used as a loading control for the evaluation of total protein levels. Results were further normalized to respective control conditions to allow for comparison between separate experiments. Following primary antibodies were used in the experiments: mouse anti-Actin (1:10000, Millipore, MAB1501), mouse anti-GAPDH (1:5000, Santa Cruz sc-322333) rabbit anti-Gsk3β (1:500, Cell Signal Technology 9315, Danvers, MA), rabbit anti-Fxr1 (1:1000, Abcam 129089), rabbit ant-Fxr2 (1:500, CST #7098), rabbit anti-Fmr1 (1:500, Abcam 17722), mouse anti-GFP (1:1000, Rockland/VWR 600-301-215), mouse anti-GluA1 (1:1000, Millipore MAB2263), mouse anti-GluA2 (1:1000, Millipore MAB397), rabbit anti-p845GluA1 (1:1000, Millipore 06.773), rabbit anti-p880GluA2 (1:1000, Abcam ab52180). Secondary antibodies: goat anti-mouse IR Dye 680 (1:10000, Mandel 926-68020), goat anti-rabbit IR Dye 800 (1:10000, Mandel 926-32211).

### Sleep deprivation

Mice were sleep deprived for 6 hours by gentle handling starting at the light onset similar to previously reported ^49, 51^. At the beginning of the sleep deprivation period, mice were taken from their home cages and transferred to a new cage and were gently handled by a researcher (e.g. using a brush) every time they were falling asleep. In the context of EEG recording, mice were kepth in the same cage for the whole procedure, including SD. At the beginning of the fourth hour, mice were transferred to a new cage again. Sleep deprivation continued until the end of the sixth hour. Littermate mice remained undisturbed in their home cages to serve as a control. Then sleep-deprived and control mice were killed by rapid cervical dislocation followed by electrophysiology recordings or Western blot experiments.

### Immunoprecipitation of polyribosomes and RNA isolation

Immunoprecipitation of polyribosomes was performed as described before ^32^. Tissue samples were lysed in homogenization buffer (50mMTris, pH 7.5, 100 mM KCl, 12 mM MgCl2, 1% Nonidet P-40, 1 mM DTT, 100U/mL RNase Out, 100 µg/mL cycloheximide, Sigma protease inhibitor mixture) followed by centrifugation for 10 min at 10000g. Anti-hemagglutinin (HA) antibody (1:150; MMS-101R; BioLegend) was added into collected supernatant and tubes were kept under constant rotation for 4 hours at 4°C. Protein G magnetic beads (Life Technologies) were washed 3 times with homogenization buffer then added into the mixture and kept for constant rotation overnight at 4°C. The following day magnetic beads were washed three times with high salt buffer (50mMTris, pH 7.5, 300 mM KCl, 12 mM MgCl2, 1% Nonidet P-40, 1 mM DTT, 100U/mL RNase Out, 100 µg/mL cycloheximide, Sigma protease inhibitor mixture). RNA was extracted by adding TRI reagent (Zymo research) to magnetic beads pellet followed by Direct-zol RNA kit according to the manufacturer’s instructions (Zymo Research). The RNA concentration was quantified using ND-1000 Spectrophotometer (NanoDrop Technologies).

### RT-PCR qPCR

RNA was extracted from neuronal cultures using TRI reagent (Zymo research) followed by Direct-zol RNA kit according to the manufacturer’s instructions (Zymo Research). Complementary DNA was synthesized using a reverse transcriptase SuperScript III kit according to the manufacturer’s instructions (Invitrogen). qPCR was performed using TaqMan™ Gene Expression Assays (Applied Biosystems) and TaqMan™ probes for Fxr1 (Thermo Fisher Scientific Mm00484523_m1), Fxr2 (Thermo Fisher Scientific Mm00839957_m1), Fmr1 (Thermo Fisher Scientific Mm01339582_m1) and Gapdh (Thermo Fisher Scientific Mm99999915_g1). Data was acquired by QuantStudio 3 Real-Time PCR System (Thermo Fisher Scientific). Relative expression analysis was performed using data from biological triplicates of each sample by QuantStudio TM Design and Analysis Software (Thermo Fisher Scientific).

### RNAseq analysis

#### Quality control

The quality control metrics for the RNA-seq data were obtained using the tool RNA-SeQC (v1.1.7). For more information, please visit their website found here: http://www.broadinstitute.org/cancer/cga/rna-seqc. This program takes aligned files as input and delivers a series of plots and statistics for each sample. Based on the output for each sample, the RNA sequencing quality was deemed acceptable for further analysis.

##### Processing Pipeline

All raw FASTQ files were aligned to the appropriate mouse genome (GRCm38) using the HISAT2 aligner. HISAT2 is a fast and sensitive alignment program that uses a large set of small graph FM (GFM) indexes that collectively cover the whole reference genome. These local indexes, in conjunction with a series of alignment strategies, ensure a rapid and accurate alignment of sequencing reads. Accessory programs for the alignment stage include SAMTOOLS (v1.3.1) and BEDTOOLS (v2.26.0). Alignment files were sorted by their genomic location and indexed using SAMTOOLS. These sorted binary SAM (BAM) files were then used as input for StringTie (v1.3.4), which assembles RNA-seq alignments into potential transcripts. It uses a novel network flow algorithm as well as an optional de novo assembly step to assemble and quantitate full-length transcripts representing multiple splice variants for each gene locus. Finally, in order to identify differentially expressed genes between samples, the Ballgown R-package was implemented (v3.4.3). Transcript-level FPKMs were estimated using Tablemaker. Expression was estimated for each transcript, exon, and intron (junction) in the assembly. All of the statistical analysis (organization, visualization, etc.) was conducted with the tools available within the Ballgown package.

### Differential Expression Analysis

The statistical test applied to this data was a parametric F-test comparing nested linear models; details are available in the Ballgown manuscript. Briefly, two models are fit to each feature, using the expression as the outcome: one including the covariate of interest (e.g., case/control status) and one not including that covariate. An F statistic and p-value are calculated using the fits of the two models. A significant p-value means that the model including the covariate of interest fits significantly better than the model without that covariate, indicating differential expression. All the differentially expressed transcripts (DETs) with p < 0.05 were selected for further analysis. Differential expression testing was carried out for the following comparisons: Ctrl/S vs Ctrl/SD and Ctrl/SD vs Fxr1over/SD.

### Gene set enrichment analysis

First DETs were filtered. All selected transcripts had a mean expression >0.5 FKPM. Fold change (FC) threshold was set to 0.7<FC>1.3. Enrichment analyses and visualization were performed following the pipeline described in Reimand et al ^34^. Enrichment analysis was performed using gProfiler (https://biit.cs.ut.ee/gprofiler/gost). Term size for pathways was selected to be min 5 and max 150. Only pathways passing significance threshold of 0.05 were selected. Gene enrichment in Gene Ontology Biological pathways (GO:BP) were selected. Then .GEM and .GMT files were downloaded from gProfiler and were used in EnrichmentMap app of Cytoscape (Ver 3.7.1) for visualization. Following parameters of EnrichmentMap were used: FDR q value cutoff <0.05, Jaccard combined > 0.375, overlap > 0.5 and Prefuse Force Directed layout was chosen. Then ClusterMaker2 App was used to cluster enriched pathways based on similarity and AutoAnnotate and WordCloud Apps were used to name clusters of enriched pathways using default parameters. The final pictures of the clusters of enriched pathways (In case of Ctrl/S vs Ctrl/SD comparison only top 10 biggest clusters are shown) are shown in Fig. S7. The list of all the enriched pathways are shown in Table S4 and S5.

Enrichment in SynGO Localization and function was performed using default parameters (https://syngoportal.org/) against brain expressed background. Significant enrichment was considered at 5% FDR (FDR<0.05). The graphical representation is shown in Fig. 4 and the whole list of pathways and genes are shown in Table S6 and S7.

### Quantification and statistical analysis

The data are presented as means ± SEM. For comparison between two groups two-tailed t-test were used. For comparison between multiple groups one-way ANOVA was used followed by Bonferroni-corrected pair-wise comparisons using GraphPadPrism 5 software (La Jolla, CA) (*p < 0.05, **p < 0.01, ***p < 0.001).

