## Supplementary Material for "Fxr1 regulates sleep and synaptic homeostasis"

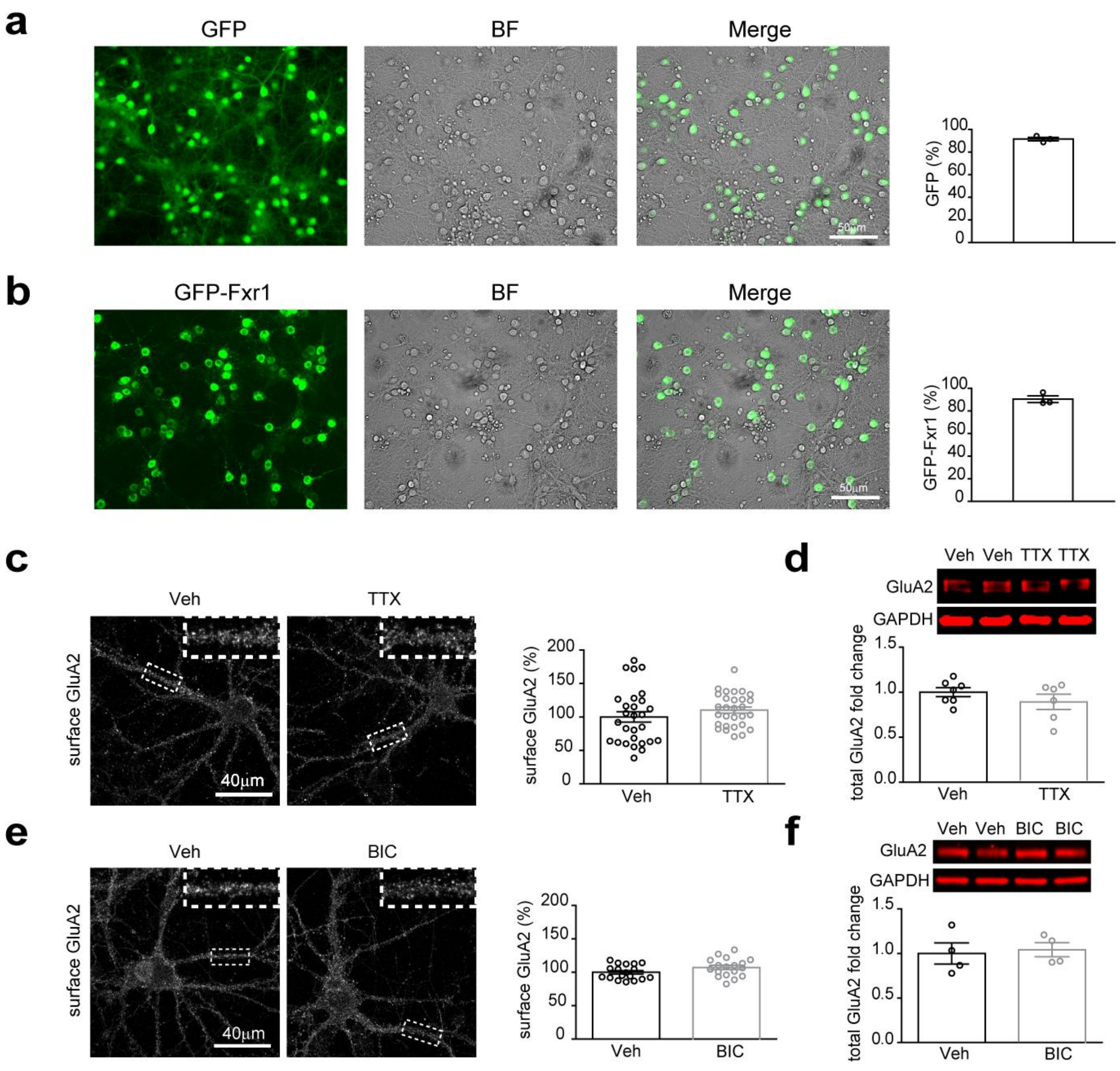


Supplementary Fig. 1

**Supplementary Fig. 1. Expression of GluA2 during synaptic scaling.**

**a** and **b**, Quantification of the % of infection by **a,** AAV1 Syn GFP or **b**, AAV1 Syn GFP-Fxr1. **c**, Surface expression of GluA2 during upscaling. **d**, Expression of total GluA2 during upscaling. **e**, Surface expression of GluA2 during downscaling. **f**, Expression of total GluA2 during downscaling. Error bars are mean ± SEM.

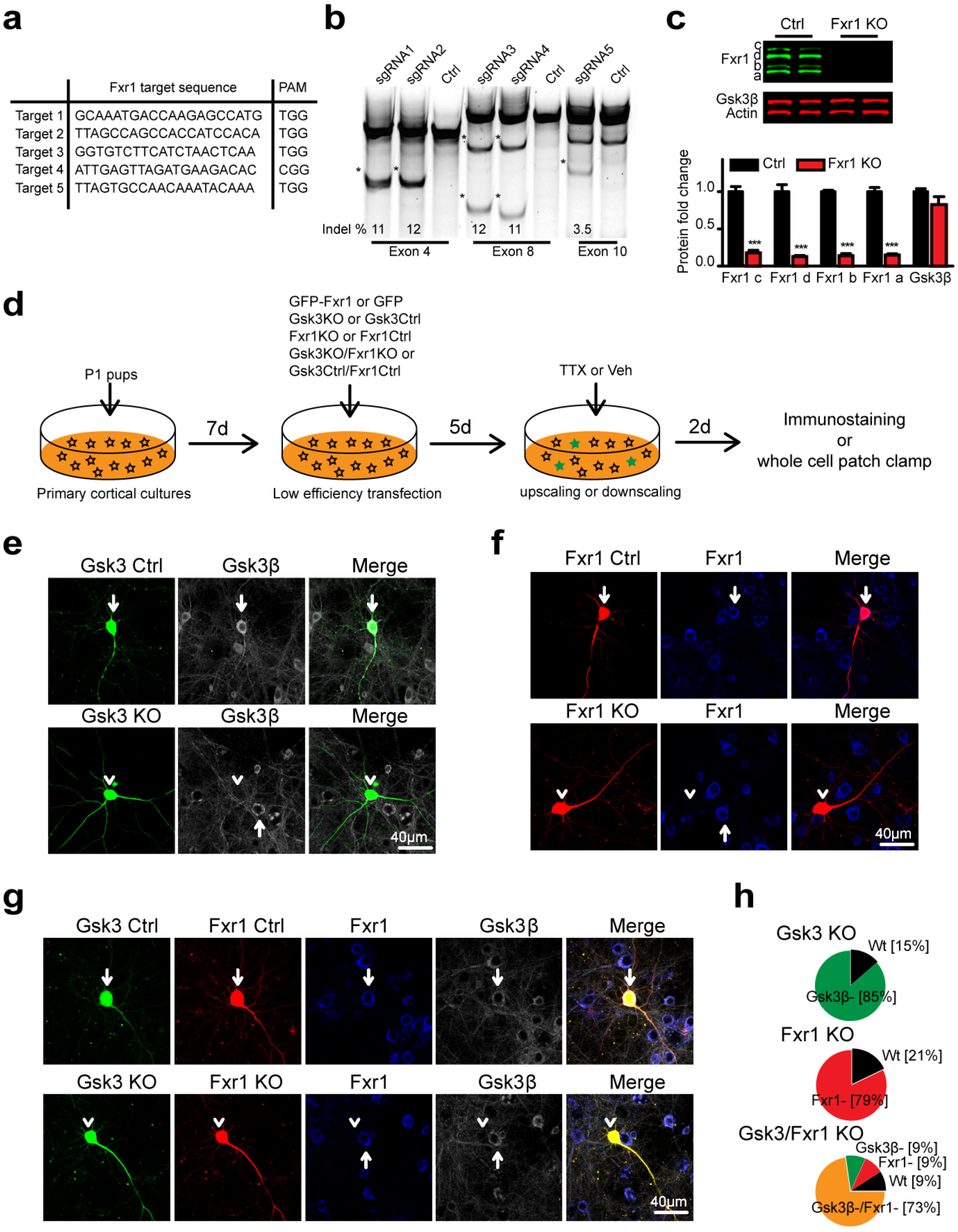


Supplementary Fig. 2

**Supplementary Fig. 2. CRISPR/Cas9 mediated knockout of *Gsk3b* and *Fxr1* in primary neurons.**

**a**, *Fxr1* targeting gRNA sequences and corresponding protospacer adjacent motifs (PAMs). **b**, Evaluation of *Fxr1* targeting sgRNAs by SURVEYOR assay 2 days after transfection of sgRNAs and SpCas9. **c**, Western blot analysis and quantification of Gsk3β and Fxr1 expression in Neuro2A cells 7 days after transfection of CRISPR/Cas9 constructs. Student's T-Test, ***$p<0.001$. Error bars are mean ± SEM. **d**, Schematic representation of low-efficiency transfection of primary neuronal cultures with various plasmids. **e-g**, Evaluation of CRISPR/Cas9 KO of **e**, *Gsk3b*, **f**, *Fxr1* and **g**, *Gsk3b/Fxr1* in primary neuronal cultures by immunostaining. Arrows indicate presence and arrowheads absence of staining. **h**, Quantification of CRISPR/Cas9 KO of *Gsk3b* and *Fxr1*.

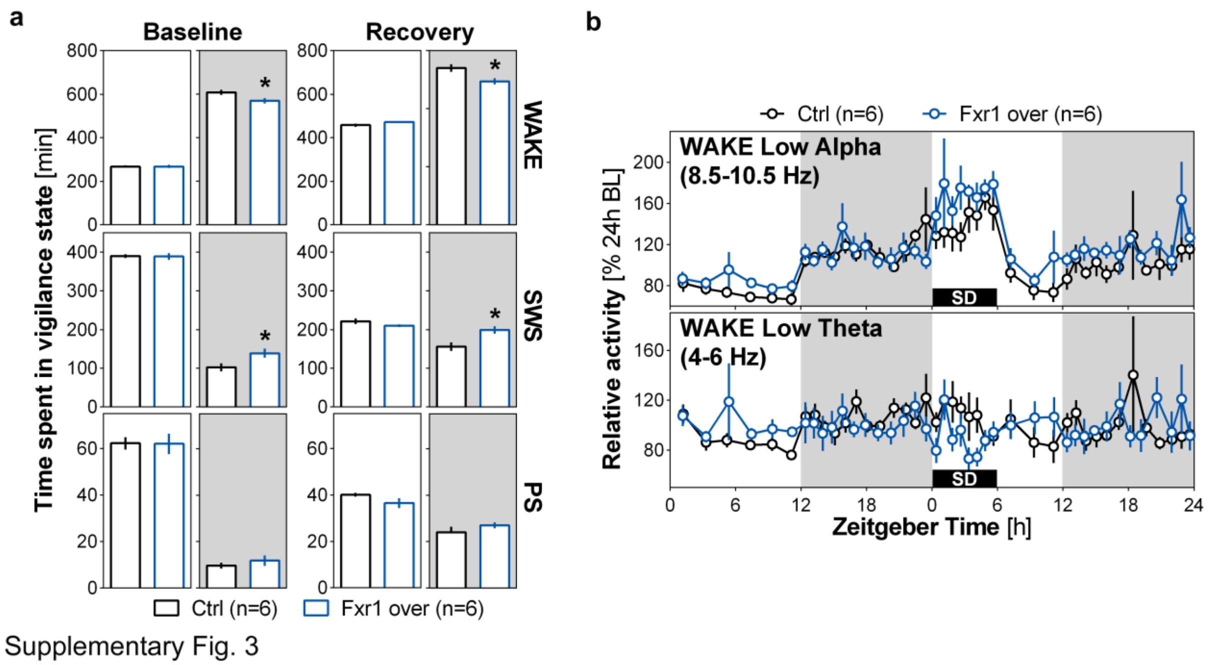


**Supplementary Fig. 3. Wakefulness and sleep duration and time course of EEG spectral activity**

**a**, Time spent in wakefulness (WAKE), slow-wave sleep (SWS) and paradoxical sleep (PS) during the 12-h light period and the 12-h dark period of the baseline (BL) recording and of the following day starting with a 6-h sleep deprivation (SD) (recovery: REC) in Ctrl or Fxr1over mice. Significant differences between groups were found for wakefulness during the 12-h dark period of both BL ($t = 2.25$) and REC ($t = 2.76$), and for SWS during the 12-h dark period of both BL ($t = 2.32$) and REC ($t = 2.93$). Dark backgrounds indicate 12-h dark periods **b**, Time course of wakefulness low alpha and low theta activity during BL and REC in Ctrl and Fxr1over mice.

**Supplementary Table 1. Differentially expressed transcripts (DETs) from Ctrl/S vs Ctrl/SD comparison.**

List of transcripts that are differentially expressed between Ctrl Sleep (Ctrl/S) and Ctrl sleep deprivation (Ctrl/SD) conditions, $p<0.05$. The table shows FKPM values for 3 individual samples in each condition.

**Supplementary Table 2. Differentially expressed transcripts from Ctrl/SD vs Fxr1over/SD comparison.**

List of transcripts that are differentially expressed between Ctrl Sleep deprivation (Ctrl/SD) and Fxr1over sleep deprivation (Fxr1over/SD) conditions, $p<0.05$. The table shows FKPM values for 3 individual samples in each condition.

**Supplementary Table 3. Common affected transcripts between Ctrl/S vs Ctrl/SD and Ctrl/SD vs Fxr1over/SD comparisons.**

List of differentially expressed transcripts that are common between Ctrl/S vs Ctrl/SD and Ctrl/SD vs Fxr1over/SD comparisons. The table shows FKPM values for 3 individual samples in each condition.

**Supplementary Table 4. List of GO:BP enriched pathways for DETs from Ctrl/S vs Ctrl/SD comparison.**

Enrichment of differentially expressed transcripts from Ctrl/S vs Ctrl/SD comparison into GO:BP performed by gProfiler. The list indicates GO:BP IDs, descriptions, enrichment FDR values, and names of genes enriched in each pathway.

**Supplementary Table 5. List of GO:BP enriched pathways for commonly affected transcripts between Ctrl/S vs Ctrl/SD and Ctrl/SD vs Fxr1over/SD comparisons.**

Enrichment of differentially expressed transcripts that are common between Ctrl/S vs Ctrl/SD and Ctrl/SD vs Fxr1over/SD comparisons into GO:BP is performed by gProfiler. The list indicates GO:BP IDs, descriptions, enrichment FDR values, and names of genes enriched in each pathway.

**Supplementary Table 6. List of SynGO:CC and SynGO:BP enrichment for DETs from Ctrl/S vs Ctrl/SD comparison.**

Enrichment of differentially expressed transcripts from Ctrl/S vs Ctrl/SD comparison into SynGO:CC and SynGO:BP is performed by SynGO online tool (https://syngoportal.org/). The list indicates SynGO:CC and SynGO:BP IDs, descriptions, enrichment FDR values, and names of genes enriched in each pathway.

**Supplementary Table 7. List of SynGO:CC and SynGO:BP enrichment for common affected transcripts between Ctrl/S vs Ctrl/SD and Ctrl/SD vs Fxr1over/SD comparisons.**

Enrichment of differentially expressed transcripts that are common between Ctrl/S vs Ctrl/SD and Ctrl/SD vs Fxr1over/SD comparisons into SynGO:CC and SynGO:BP is performed by SynGO online tool (https://syngoportal.org/). The list indicates SynGO:CC and SynGO:BP IDs, descriptions, enrichment FDR values, and names of genes enriched in each pathway.
